# *Fusarium verticillioides* FvPex8 is a key component of peroxisomal docking/translocation module that serves important roles in fumonisin biosynthesis but not in virulence

**DOI:** 10.1101/2020.05.06.081687

**Authors:** Wenying Yu, Mei Lin, Minghui Peng, Huijuan Yan, Jie Zhou, Guodong Lu, Won Bo Shim, Zonghua Wang

## Abstract

Peroxisomes are ubiquitous organelles in eukaryotic cells that fulfill various important metabolic functions. In this study, we investigated the role of Docking/Translocation Module (DTM) peroxins, mainly FvPex8, FvPex13, FvPex14, and FvPex33, in *Fusarium verticillioides* virulence and fumonisin B1 (FB1) biosynthesis. Protein interaction experiments suggested that FvPex13 serves as the core subunit of *F. verticillioides* DTM. When we generated gene deletion mutants (ΔFvpex8, ΔFvpex13, ΔFvpex14, ΔFvpex33, ΔFvpex33/14) and examined whether the expression of other peroxin genes were affected in the DTM mutants, ΔFvpex8 strain showed most drastic changes to *PEX* gene expression profiles. Deletion mutants exhibited disparity in carbon source utilization and defect in cell wall integrity when stress agents were applied. Under nutrient starvation, mutants also showed higher levels of lipid droplet accumulation. Notably, ΔFvpex8 mutant showed significant FB1 reduction and altered expression of *FUM1* and *FUM19* genes. However, FvPex13 was primarily responsible for virulence, while ΔFvpex33/14 double mutant also showed virulence defect. In summary, our study suggests that FvPex13 is the core component of DTM, regulating peroxisome membrane biogenesis as well as PTS1- and PTS2-mediated transmembrane cargo transportation. Importantly, we predict FvPex8 as a key component in DTM that affects peroxisome function in FB1 biosynthesis in *F. verticillioides*.

## Introduction

Peroxisomes are a family of membrane-bound microbodies found in all major eukaryotes, with specific metabolic properties showing substantial differences among various species and tissue types (Alberts *et al*., 2002, Cooper, 2000). These cellular organelles, including peroxisomes, glyoxysomes in plants and fungi, glycosomes in trypanosomes, and Woronin bodies in filamentous fungi, are involved in a variety of important cellular functions, including fatty acid metabolism, reactive oxygen species detoxification, and secondary metabolite biosynthesis (Saikia & Scott, 2009, Imazaki *et al*., 2010, Maggio-Hall *et al*., 2005, Reverberi *et al*., 2012, Chen *et al*., 2018). Dysfunctional peroxisomes have been shown to result in severe and lethal disorders in humans and mammals (Cho *et al*., 2018, Dixit *et al*., 2010, Thoms *et al*., 2009, You *et al*., 2015). Plant glyoxysomes play essential roles in glyoxylate cycle, embryonic development, and gametophytic function (Hu *et al*., 2012, Boisson-Dernier *et al*., 2008). Glycosomes in Trypanosomes perform important glycolytic reactions (Bauer & Morris, 2017). In filamentous fungi, a specific type of peroxisome known as Woronin bodies are associated with septal pore maintenance and cellular integrity (Jedd & Chua, 2000). Recent studies also suggest that peroxisomes play important roles in development, secondary metabolism, and virulence (Kimura *et al*., 2001, Wang *et al*., 2013, Soundararajan *et al*., 2004), but underlying genetic mechanisms associated with these fungal-specific functions still warrant further investigation.

*Fusarium verticillioides* is a major pathogen of maize, particularly due to ear rot disease and mycotoxin contamination worldwide (Marin *et al*., 2013, Wu *et al*., 2014, Woloshuk & Shim, 2013). It is also known as the pathogen responsible for pokkah boeng in sugarcane, and approximately 90% of this disease in China is attributed to *F. verticillioides* (Zhang *et al*., 2015, Hilton *et al*., 2017, Lin *et al*., 2014). Interestingly, Zhang et al (Zhang *et al*., 2018b) recently discovered a key peroxisomal matrix protein (FVEG_11334, which is designated as FvPex33 in this study) and a Woronin body associated protein (FvHex1) interacting with Fsr1, a striatin homolog that regulates virulence in *F. verticillioides*. However, gene mutagenesis experiment revealed that FvPex33 and FvHex1 do not directly influence *F. verticillioides* virulence when tested on maize seedlings. This outcome, along with earlier published studies on the role of peroxins in fungal virulence and secondary metabolism (Fujihara *et al*., 2010, Min *et al*., 2012, Li *et al*., 2017) suggested that other peroxisomal matrix proteins may play important roles in *F. verticillioides* virulence and mycotoxin biosynthesis.

Biogenesis of peroxisomes is dynamically regulated in response to environmental and developmental cues (Lazarow, 2003, Heiland & Erdmann, 2005). Metabolic tasks and enzyme content are adjusted according to taxa-specific cellular needs through generation and degradation (Pieuchot & Jedd, 2012, Nuttall *et al*., 2011). There are two primary pathways for peroxisome formation: *de novo* biogenesis and mature peroxisomes fission. The *de novo* synthesis of peroxisomes has two steps: formation of the peroxisomal membrane and subsequent sorting of peroxisomal matrix proteins to the organelle (Opalinski *et al*., 2011). The peroxisomal importomer model helps to explain how newly synthesized matrix proteins bind to the cytosolic receptor at the peroxisome membrane and form a highly dynamic pore to transfer matrix proteins to peroxisome lumen (Meinecke *et al*., 2010). These transient cargo-docking-translocation complexes are formed by variable combination of peroxin (Pex) proteins (Table S1). Briefly, the key docking complex core consists of the SH3-domain protein Pex13 and the PXXP-motif protein Pex14, but also harbors additional factors, e.g. Pex17 in lower eukaryotic cells, Pex33 (also known as Pex14/17) in filamentous fungi, and Pex13.2 in *Trypanosoma brucei* (Verplaetse *et al*., 2012). A recent study also characterized Pex17/Pex14/Dyn2p complex in yeast (Chan *et al*., 2016). Physical association among docking peroxins varies in different organisms, and recent reviews extensively explain how various matrix proteins with type 1 and type 2 peroxisomal targeting signal (PTS1 and PTS2) are recognized by the specific docking complex and associate with the peroxisomal membrane (Nuttall *et al*., 2011, Pieuchot & Jedd, 2012).

PTS1 and PTS2 peroxisomal matrix proteins are recognized by specific receptors Pex5 and Pex7, respectively, and recruited to the peroxisome docking complex (Meinecke *et al*., 2010, Heiland & Erdmann, 2005). In addition to these core Peroxins in docking/translocation module (DTM), other Pex proteins are dynamically involved in translocating matrix proteins across the peroxisome membrane (Platta *et al*., 2013). In particular, Pex8 is known to be associated with DTM but is only present in yeasts and filamentous fungi (Agne *et al*., 2003). It has been suggested that Pex8 can serve as a importomer-bridging protein to hold the docking complex and the export machinery complex together to facilitate dynamic and transient translocation (Rayapuram & Subramani, 2006, Platta *et al*., 2013, Meinecke *et al*., 2010), but the detailed function is largely unknown. In this study, our aim was to characterize putative *F. verticillioides* peroxins, particularly those predicted to be involved in DTM. While paying a close attention to two core components of the docking complex, Pex13 and Pex14, we expanded our study into additional peroxisomal proteins, namely Pex8, that have not been well characterized in plant pathogenic fungi. Through a series of fungal metabolism, physiology and virulence assays, we concluded that Pex8, along with Pex13, plays a critical role in the generation of docking complex in peroxisomes and for peroxisome function in *F. verticillioides* virulence and mycotoxin biosynthesis.

## Results

### Protein-protein interactions between *F. verticillioides* DTM components and other key peroxins

Peroxins Pex13 and Pex14 are recognized as key peroxisomal membrane docking complex proteins in eukaryotes, which are required for protein import into peroxisomes. In addition, Pex33 has been identified as a new component of the docking complex in *Neurospora crassa* (Managadze *et al*., 2010), and Pex8 is known to be associated with DTM in yeasts and filamentous fungi (Agne *et al*., 2003). Using these proteins as query, we screened for putative DTM components, *i.e*. Pex13, Pex14, Pex33, and Pex8, in *F. verticillioides* using the BLASTP algorithm. Additional details for these key Pex proteins, e.g. sequence similarities and putative functional motifs, are provided in Table S1 and Fig. S1A. Notably, further comparison of FvPex33 against Pex14- and Pex17-like proteins in *S. cerevisiae, F. graminearum, F. verticillioides* strongly suggested that FvPex33 is distinct from Pex14 and Pex17 and is a unique component of DTM in *F. verticillioides* (Fig. S1B).

Based on Prediction of Transmembrane Regions and Orientations (TMpred) algorithm (Hoffman and Stoffel, 1993), FvPex8, FvPex13, FvPex14, and FvPex33 are predicted to have transmembrane helices and likely belong to peroxisomes membrane proteins (Fig. S1C). To study the cellular localization of these proteins in *F. verticillioides*, we first chose the biogenesis protein Pex14 as the peroxisomal marker and prepared Pex14-mCherry construct (Grant *et al*., 2013). We studied transformants containing GFP-fused Pex8, Pex13, or Pex33 protein along with Pex14-mCherry for overlapping red and green fluorescence signals (Fig. 1A), and the images revealed Pex14 co-localizing with Pex13 and Pex33 while showing partial overlap with Pex8.

**Fig. 1.**
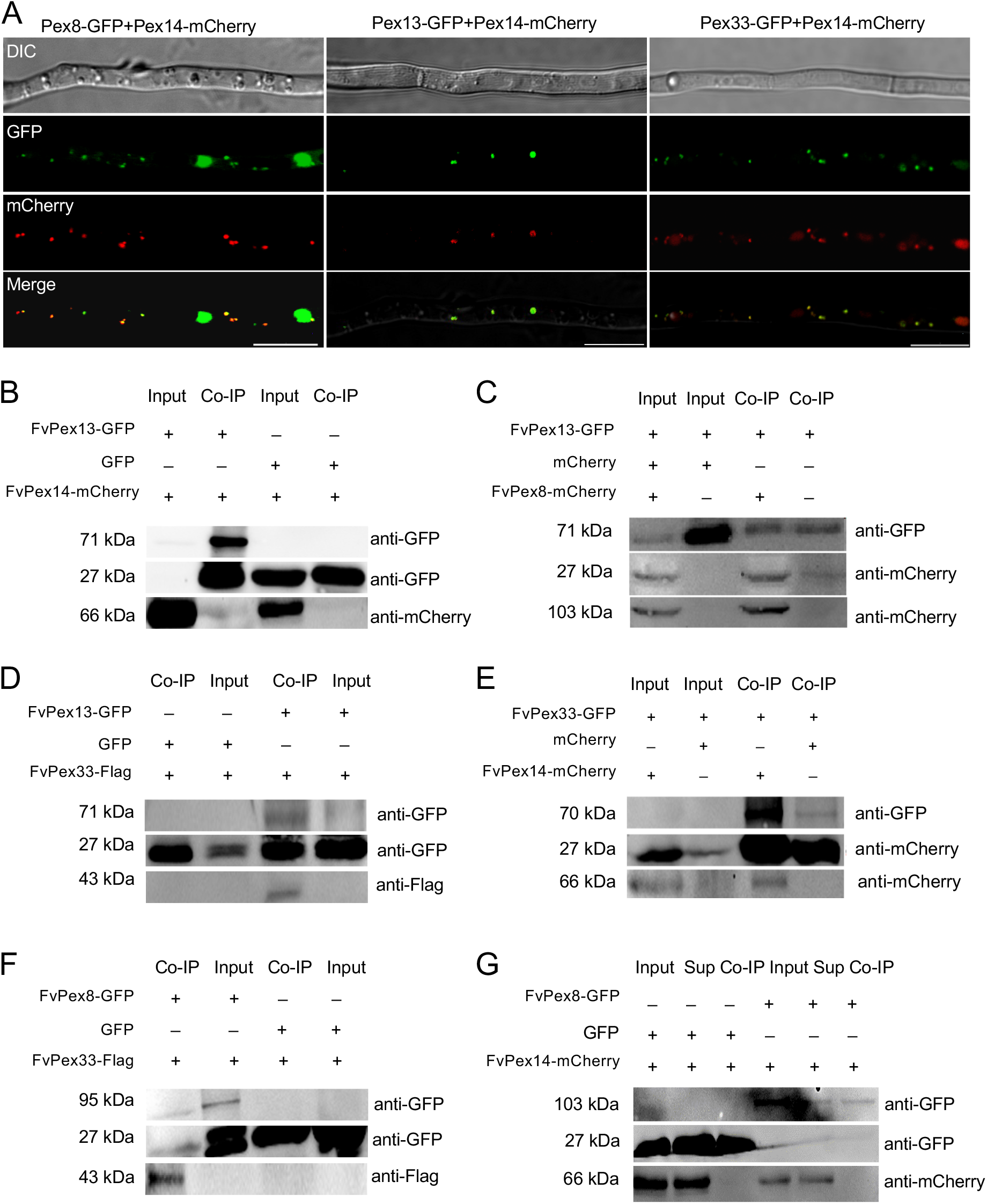
FvPex8, FvPex13, FvPex14, and FvPex33 protein interaction analyses. (A) Localization of FvPex8-GFP, FvPex13-GFP, and FvPex33-GFP were studied in comparison to FvPex14-mCherry, which was used as a peroxisomal marker, in the wild-type strain. Scale bar = 10 μm. GFP-trap-based pull-down experiment showing the interaction *in vitro* between (B) FvPex13 and FvPex14, (C) FvPex13 and FvPex8, (D) FvPex33 and FvPex13, (E) FvPex33 and FvPex14, (F) FvPex33 and FvPex8, and (G) FvPex8 and FvPex14. The strains co-expressing the indicated proteins were immunoprecipitated with GFP trap beads. The IP signal (FvPex13-GFP, FvPex8-GFP, or FvPex33-GFP) and the Co-IP signal (FvPex8-mCherry, FvPex14-mCherry, or FvPex33-Flag) were detected by immunoblots probed with antibodies to GFP, mCherry, and Flag, respectively.

To further verify interactions between key DTM subunits and peroxins, we co-expressed following six pairs of proteins, *i.e*. FvPex13-GFP/FvPex14-mCherry, FvPex13-GFP/FvPex8-mCherry, FvPex13-GFP/FvPex33-Flag, FvPex33-GFP/FvPex14-mCherry, FvPex8-GFP/FvPex33-Flag, and FvPex8-GFP/FvPex14-mCherry, into *F. verticillioides* for coimmunoprecipitation (Co-IP) experiments. Results showed that FvPex13-GFP successfully associated with the beads and that FvPex14 was eluted in samples that co-expressed FvPex13-GFP and FvPex14-mCherry, indicating FvPex13 and FvPex14 binding *in vivo* (Fig. 1B). Similarly, FvPex13 successfully associated with FvPex8 and FvPex33 (Fig. 1C and D). FvPex33 successfully associated with FvPex14 and FvPex8 (Fig. 1E and F), but FvPex8 did not coprecipitate with FvPex14 (Fig. 1G). We further tested these interactions by yeast two-hybrid assays (Fig. S2A), where FvPex13 showed strong positive interaction with key DTM proteins tested. FvPex8 showed strong interaction with FvPex13 but not with FvPex14. Based on these outcomes, we propose that FvPex13 serves as the core subunit in *F. verticillioides* DTM.

### Vegetative growth and conidiation are differentially affected by carbon sources in DTM peroxin mutants

To study the function of key DTM peroxins in *F. verticillioides*, we generated ΔFvpex8, ΔFvpex13, ΔFvpex14, and ΔFvpex33 through targeted gene replacement with the hygromycin-resistance marker in *F. verticillioides* 7600 wild-type strain as described previously (Zhang *et al*., 2018b). We also generated ΔFvPex33/14 double mutant by replacing *FvPEX14* gene with the geneticin-resistance marker in ΔFvpex33 protoplast. Subsequently, we also generated complementation strains ΔFvpex8-C, ΔFvpex13-C, ΔFvpex14-C, and ΔFvpex33-C by transforming the corresponding mutant with a construct harboring the gene (with its native promoter and terminator) fused with GFP and the geneticin resistance marker. Homologous recombination was confirmed by polymerase chain reaction (PCR) the gene and quantitative PCR (qPCR) the expression of the gene in corresponding complementation strains (Supplementary Information).

To test whether these mutations impact *F. verticillioides* growth on various carbon nutrients, we cultivated these strains on minium medium (MM) and complete (CMII) agar plates with different carbon sources. On MM agar, we observed varying growth inhibition rates and colony morphologies depending on amended carbon nutrient (Fig. 2A). No strain showed particularly detrimental growth defect on MM, suggesting that these genes are not critical for vegetative growth. But on MM agar with sucrose as the carbon nutrient, vegetative growth and aerial mycelia production in ΔFvpex8 and ΔFvpex33/14 was notably hindered (Fig. 2A and 2B). Inhibition rate was more drastic in ΔFvpex8 on MM agar with butyrate and palmitic acid (Fig. 2A and 2B). Interestingly, aerial mycelium was completely absent in ΔFvpex8 mutant on media with palmitic acid. On CMII agar with sucrose as the carbon source, only ΔFvpex8 exhibted slight growth inhibition but overall all strains were morphologically very similar (Fig. 3SA and 3SB). We also observed a drastic reduction in conidiation in ΔFvpex13 and ΔFvpex33/14 mutants on CMII agar with sucrose (Fig. S3C). However, on CMII with linoleic acid growth inhibition in ΔFvpex8 was more drastic compared to other mutants (Fig. 3SA and 3SB).

**Fig. 2.**
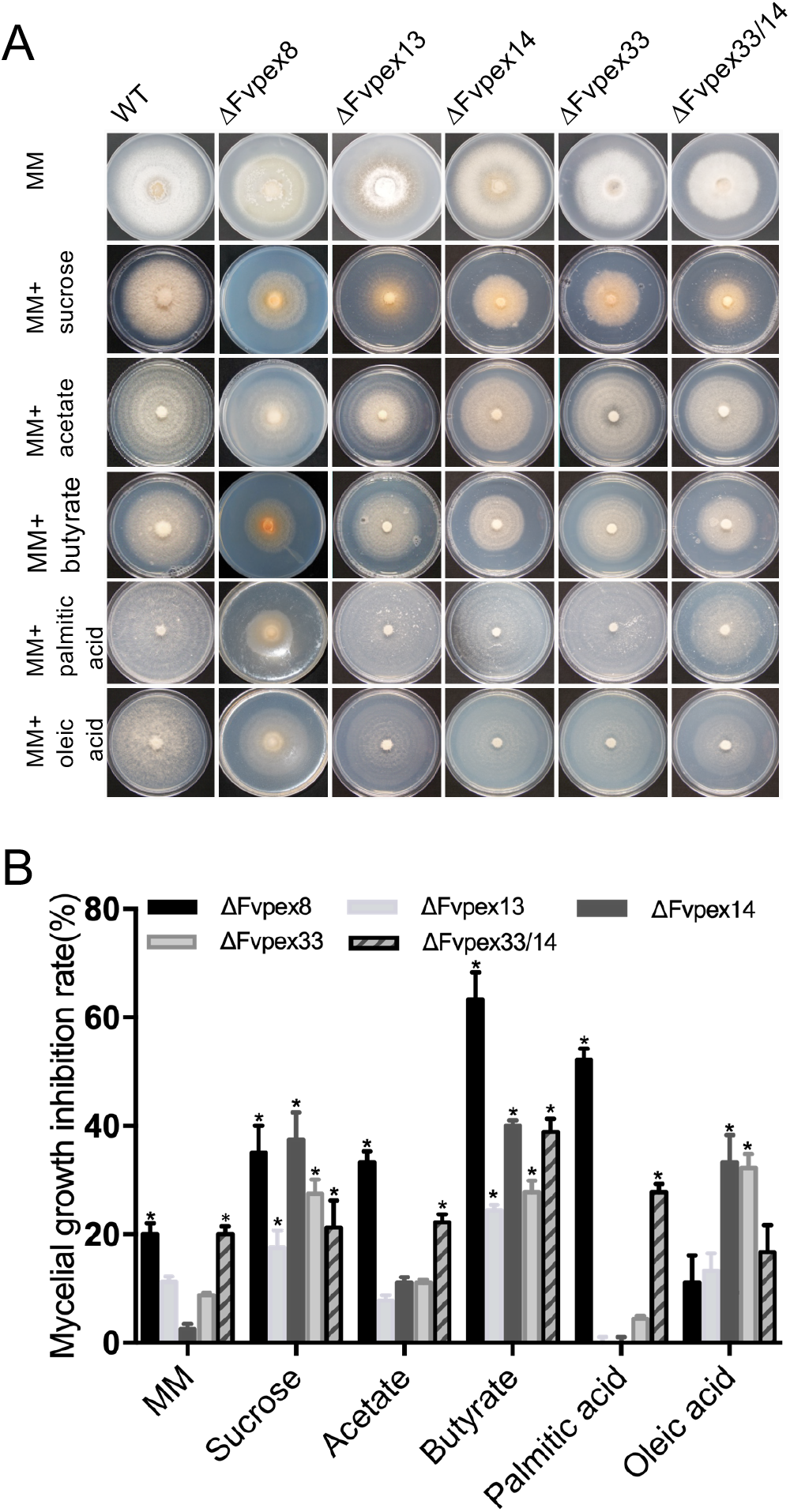
Impact of FvPex8 on *PEX* gene transcription and carbon source utilization. (A) Vegetative growth and aerial mycelia production in wild-type (WT) and mutant strains were monitored on minimal medium (MM) amended with different carbon sources. Colony morphology was photographed after 5 days of incubation at 25°C. (B) The growth inhibition rate was analyzed by comparing the growth of wild type with mutants on MM agar with various carbon amendments. The mycelial growth inhibition rate (%) was measured by [(the diameter of WT strain - the diameter of mutant strain)/(the diameter of WT strain)]×100%). Bar indicates the standard deviation of three replicates. An asterisk above the column indicates a statistically significant difference (P<0.05) as anlyzed by t-test.

### Redox homeostasis, cell wall integrity, and lipid degradation in DTM mutants

Peroxisomes serve as one of the primary sources of endogenous and exogenous ROS production. Accumulation of ROS and disruption of redox homeostasis can lead to lower cell survival (Min *et al*., 2012). When stained with CM-H2DCFDA, mutant hyphae showed green fluorescent granules indicative of high ROS accumulation, especially in ΔFvpex8, ΔFvpex13 and ΔFvpex33/14 mutants (Fig. 3A and 3B). The colonies were stained with 3,3’-diaminobenzidine (DAB) and nitro blue tetrazolium (NBT) for H_2_O_2_ and O^2^-detection, respectively. Yellow and blue colonies were observed in ΔFvpex8, ΔFvpex13, and ΔFvpex33/14 mutants with DAB and NBT staining, respectively (Fig. 3C). Peroxin mutants are likely to display cell wall integrity defects due to their inability to import enzymes of the glyoxylate cycle into peroxisomes (Bhambra et al. 2006). Almost all mutants exhibited sensitivity to calcofluor while except for ΔFvpex8 mutant (Fig. 4A). Mutants showed much higher sensitivity in general to sodium dodecyl sulfate (SDS) when compared to the wild type (Fig. 4A). We also observed growth inhibition in DTM mutants on medium amended with H_2_O_2_ when compared to the WT, which suggested that ability to respond to reactive oxygen species and maintain redox homeostasis was disturbed in these peroxin mutants (Fig. 4A). Growth inhibition rate in peroxin mutants was significantly higher than that of wild type, suggesting DTM peroxins are important for maintaining *F. verticillioides* cell wall integrity (Fig. 4B). We also studied lipid droplet degradation and turnover defects in DTM mutants. The mutants accumulated higher levels of lipid droplets compared to WT, particularly when shifted to nutrient starvation (Fig. S4).

**Fig. 3.**
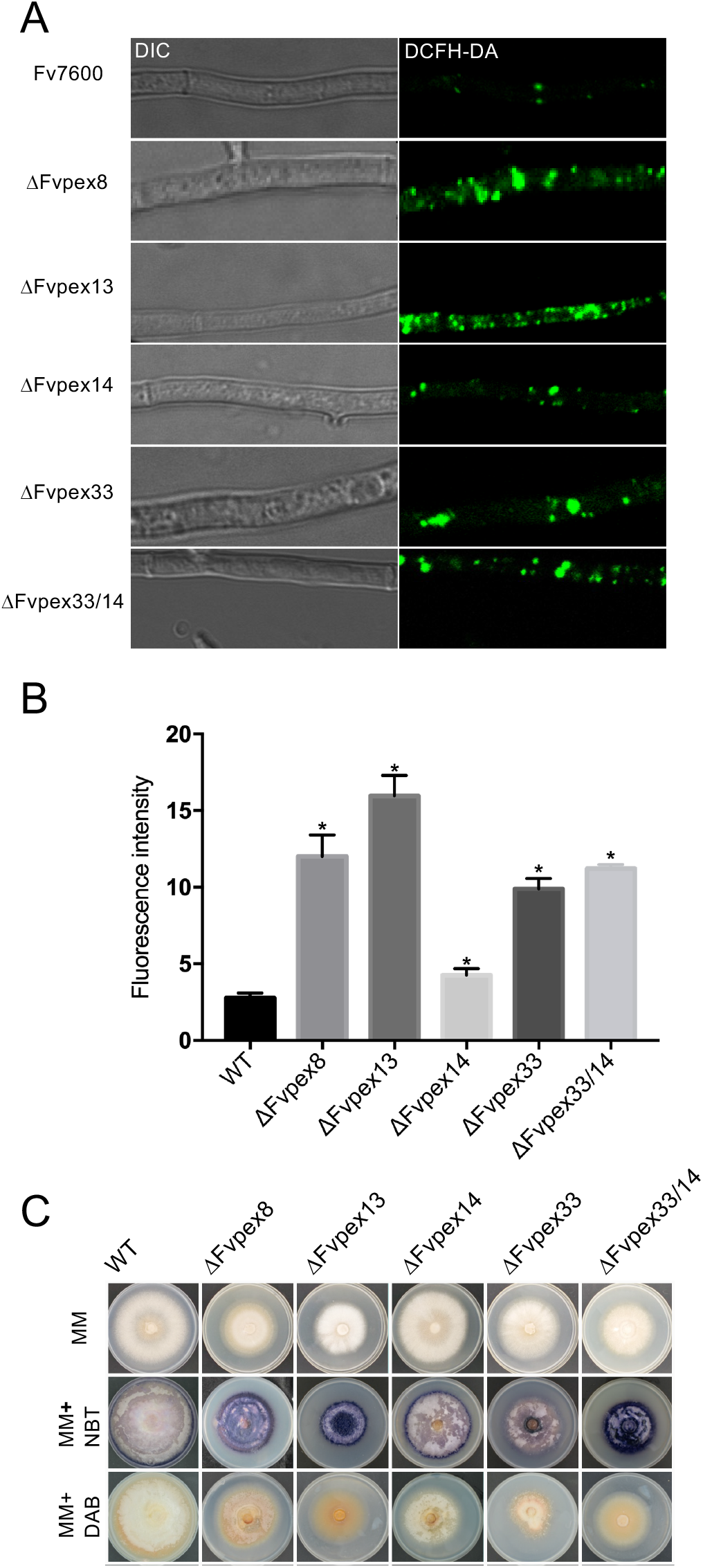
Reactive oxygen species response in mutants. (A) Fungal hyphae were grown for 2 days and stained with CM-H2DCFDA to measure cellular ROS accumulation in wild-type strain Fv7600 and mutants. Accumulation of ROS in mycelia was visualized with 6-chloromethyl-2’,7’-dichlorodihydrofluorescein diacetate (CM-H2DCFDA) staining. Scale bar = 10 μm. (B) ROS accumulation was quantified by Image J software. Error bars represent the deviation from three replicates and an asterisk above the column indicates a statistically significant difference (P<0.05) as anlyzed by t-test. (C) Endogenous ROS scavenging capacity was tested by staining fungal strains grown on CMII medium for 5 days with 3,3’-diaminobenzidine (DAB) for H_2_O_2_ detection and with Nitro Blue Tetrazolium (NBT) for O^2-^ detection.

**Fig. 4.**
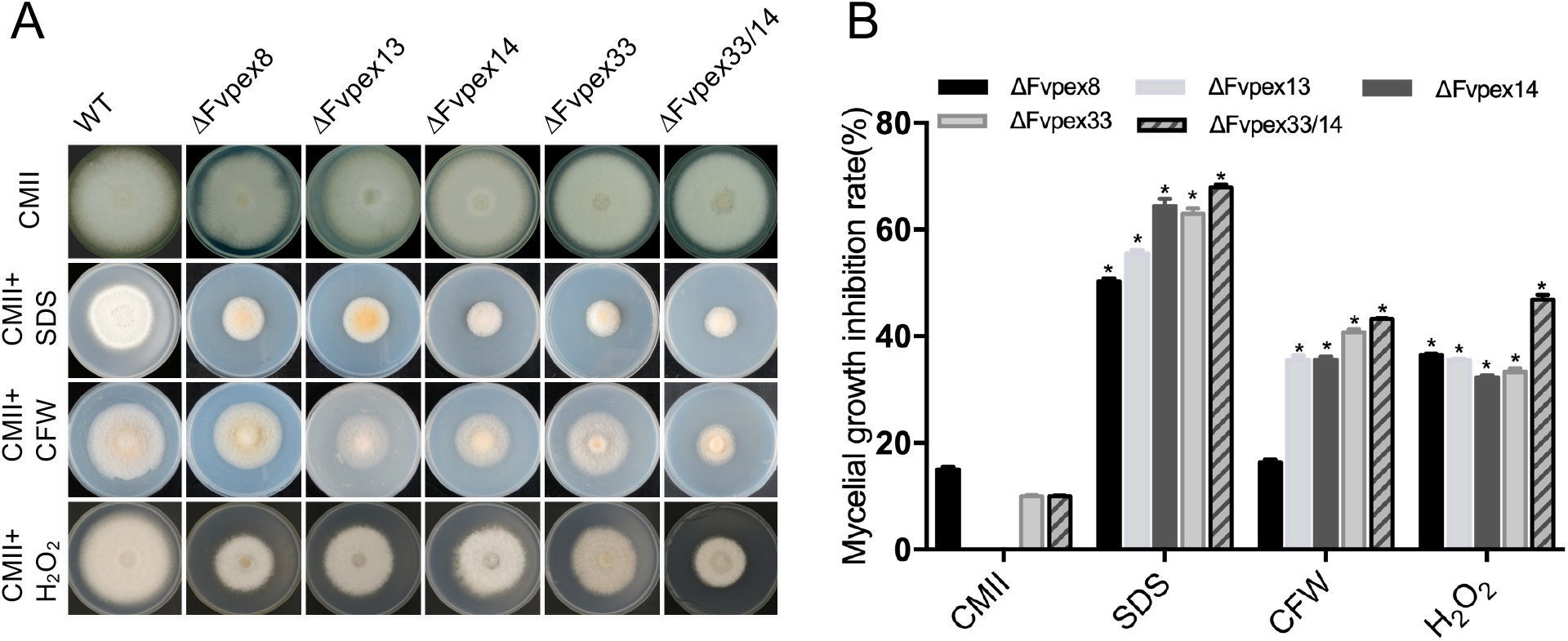
Disruption of peroxisomal functions promotes reactive oxygen species (ROS) accumulation. (A) Mycelial growth of the wild type (WT) and mutants was monitored on CMII media supplemented with sodium dodecyl sulfate (SDS), Calcofluor white (CFW), and H_2_O_2_ for 5 days. Plates were photographed after 5 days of incubation at 25°C. (B) Mycelial growth inhibition rates were calculated by [(the diameter of WT strain - the diameter of mutant strain)/(the diameter of WT strain)]×100%. An asterisk above the column indicates a statistically significant difference (P<0.05) as anlyzed by t-test.

### DTM peroxin FvPex8 function is necessary for FB1 production in *F. verticillioides*

*F. verticillioides* is one of the key fungal pathogens responsible for fumonisin contamination of maize worldwide. To assess fumonisin B1 (FB1) accumulation, we independently inoculated B73 maize kernels with spore suspensions of WT, ΔFvpex8, ΔFvpex13, ΔFvpex14, ΔFvpex33, and ΔFvpex33/14 strains. After 10 days of incubation, fungal mass in samples was assessed by qPCR quantification of β-tubulin gene *TUB2* DNA. Fungal mass in ΔFvpex8, ΔFvpex13 and ΔFvpex33/14 mutants was approximately half of those observed in WT, ΔFvpex14, and ΔFvpex33 (Data not shown). When FB1 levels were normalized to fungal mass in each sample, all mutants showed reduction in FB1 production, ranging from 50% to 94%, with ΔFvpex8 showing the most drastic reduction (Fig. 5A).

**Fig. 5.**
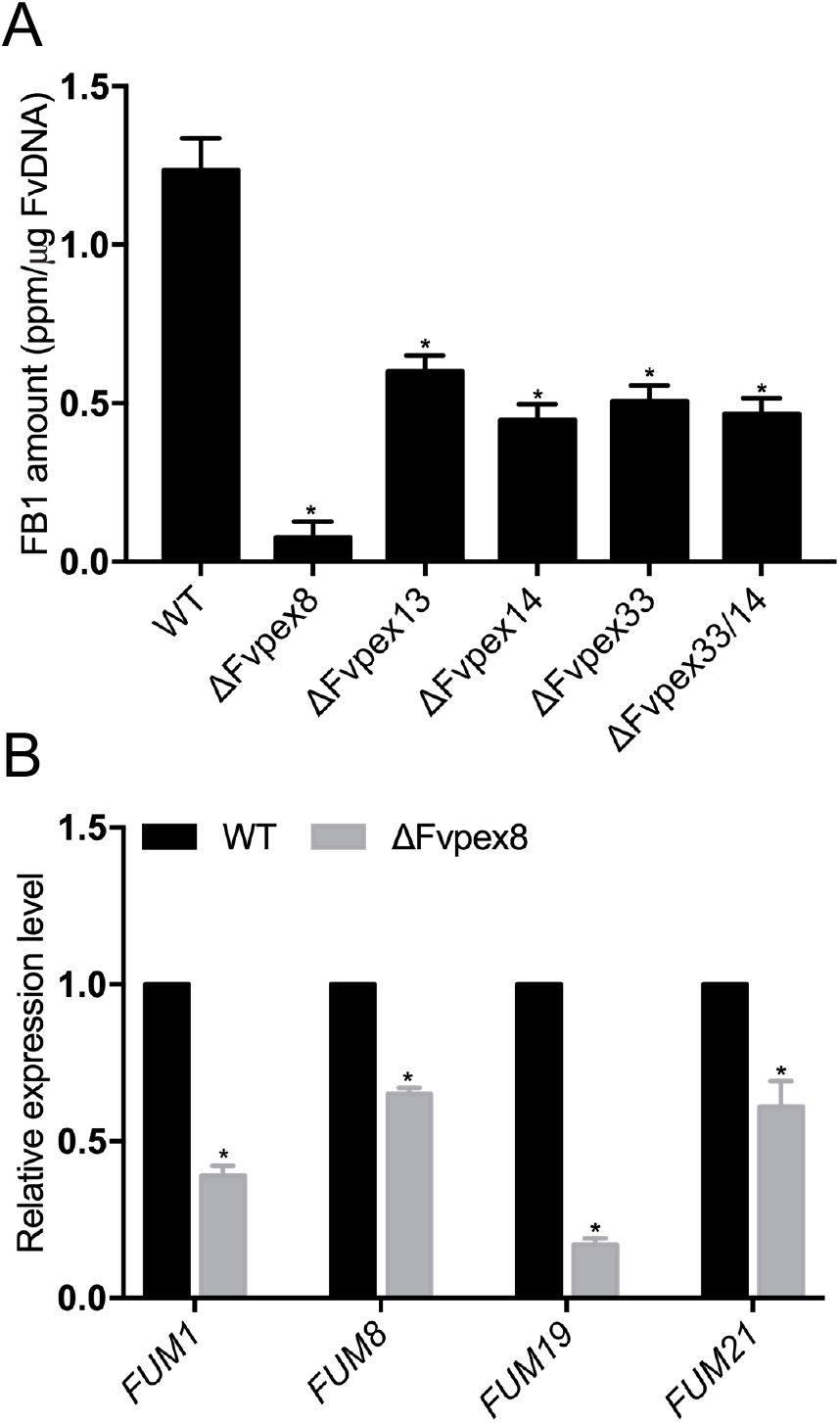
Role of peroxins in fumonisin B1 (FB1) production and *FUM* gene expression. (A) Surface sterilized B73 corn kernels were inoculated with wild type (WT) and mutant fungal conidia and were incubated for 10 days before FB1 levels were quantified by ELISA Kit. *F. verticillioides* biomass was quantified by measuring *F. verticillioides* β-tubulin (*TUB2*) DNA in samples. FB1 level in samples was normalized by calculating FB1 (ppm)/ *TUB2* DNA (ng). (B) To compare the expression levels of key *FUM* genes, RNA samples were prepared from the wild type (WT) and ΔFvpex8 mutant on corn kernel for 10 days. The expression levels of *FUM1, FUM8, FUM19* and *FUM21* were normalized to *TUB2* expression. The experiment was repeated three times, and error bars represent the standard deviation from three replicates. An asterisk above the column indicates a statistically significant difference (P<0.05) as anlyzed by t-test.

To further understand how FvPex8 mutation resulted in such drastic reduction in FB1 biosynthesis, we analyzed transcriptional expression of four key *FUM* genes: polyketide synthase *FUM1*, aminotransferase *FUM8*, transporter *FUM19*, and transcription factor *FUM21* (Bojja *et al*., 2004, Desjardins & Proctor, 2007, Brown *et al*., 2007). The expression levels of these genes were assessed by qPCR in WT and ΔFvpex8 strains after a 10-day incubation on corn kernels (Fig. 5B). When compared to the WT, the expression of these four genes were significantly suppressed in ΔFvpex8, particularly *FUM1* showing drastically lower expression levels (Fig. 5B). These outcomes suggest that FvPex8 function is necessary for proper expression of *FUM* genes in *F. verticillioides*, and ultimately fumonisin production.

### DTM peroxin FvPex13 plays key roles in *F. verticillioides* virulence

To test whether these DTM Peroxins are associated with virulence, we inoculated B73 maize stalk internodes with spore suspension of the wild-type, ΔFvpex8, ΔFvpex13, ΔFvpex14, ΔFvpex33, and ΔFvpex33/14 strains. When stalks were split open longitudinally after a 10-day incubation, ΔFvpex13 and ΔFvpex33/14 mutants showed drastically reduced rot when compared with the wild-type progenitor and other mutants (Fig. 6A and B). We further tested virulence on sugar cane (cultivar Badila), and outcomes were quite similar to those observed in maize stalk rot assays with ΔFvpex13 showing significantly decreased levels of rot (Fig. S5A and B). It is notable while ΔFvpex14 and ΔFvpex33 single mutations did not impact stalk rot, the double mutant ΔFvpex33/14 showed a significant reduction in virulence when tested on maize and sugarcane stalks. Significantly, ΔFvPex8 mutation did not have any negative impact on *F. verticillioides* virulence. FvPex13-complemented strain showed wild type-level stalk rot symptoms, suggesting that FvPex13 plays a critical role in *F. verticillioides* virulence on maize and sugarcane.

**Fig. 6.**
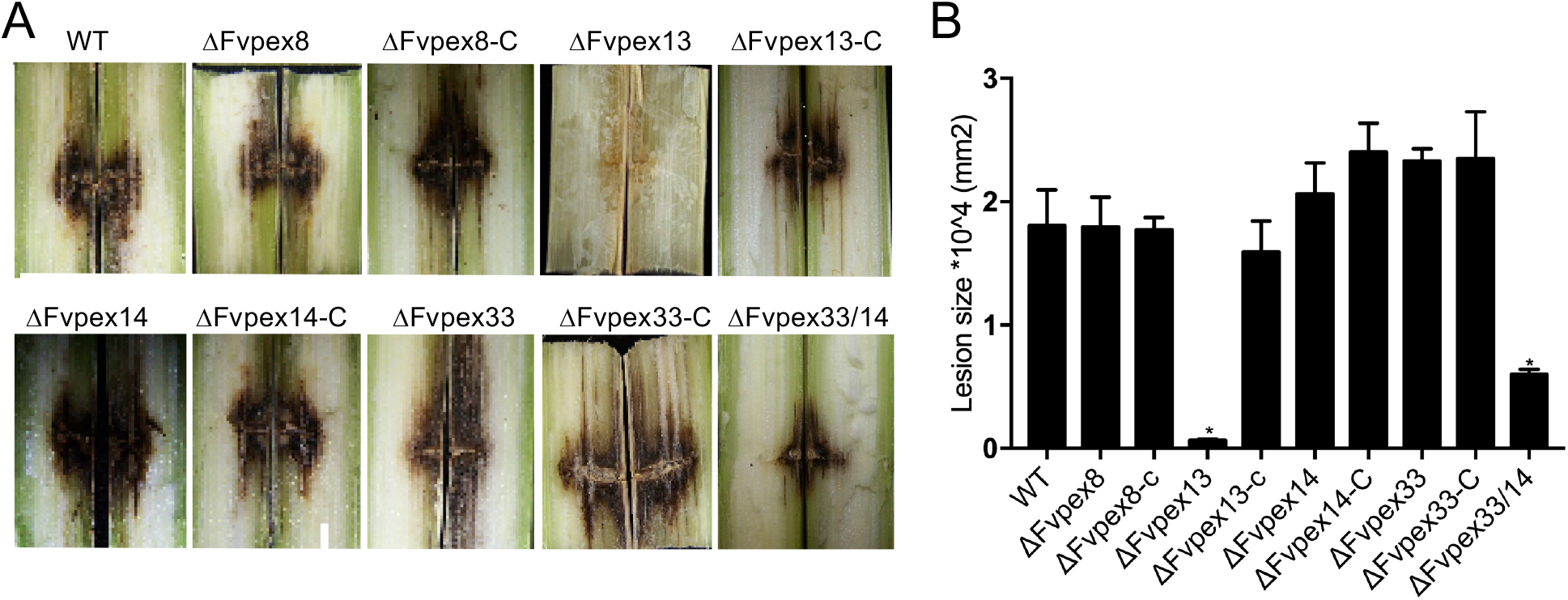
Role of DTM peroxins in maize stalk rot virulence. (A) Maize B73 stalks were inoculated with wild type (WT) and mutant fungal spore suspension (10^6^/ml) at the internodal region and incubated at room temperature for 7 days. Subsequently, maize stalks were split longitudinally to visually inspect rot symptoms. (B) Image J software was used to quantify the extent of the rot in each sample. Three independent biological repetitions were performed and error bars represent the standard deviation from three replicates. An asterisk above the column indicates a statistically significant difference (P<0.05) as anlyzed by t-test. Sterile water was used as a negative control (Fig. S5).

### Deletion of Pex13 impacts cellular localization of DTM peroxins in *F. verticillioides*

A mutation in Pex13 has been linked to PTS1 import defects in yeast, peroxisome abnormalities and appressorium defects in *Colletotrichum orbiculare* (Erdmann & Blobel, 1996, Fujihara *et al*., 2010). To monitor PTS matrix protein import in *F. verticillioides*, we used confocal microscopy to follow GFP derivatives targeted to the peroxisome. We fused GFP to carnitine O-acetyl transferase gene *FvCAT* containing C-terminal PTS1 sequence (Zolman & Bartel, 2004) or peroxisomal 3-oxoacyl-CoA thiolase gene *FvFOX3* containing *N*-terminal PTS2 sequence (Einerhand *et al*., 1995) obtaining fusion protein GFP-PTS1 and PTS2-GFP, respectively. We also tested physical interactions between these two PTS proteins and PTS receptor proteins by yeast two-hybrid assays; FvPex5 showed positive results with FvCAT and FvPex7 showed positive results with FvFOX3 (Fig. S2B and C). GFP-PTS1 showed a diffused fluorescence distributed throughout the cytoplasm while PTS2-GFP was restricted to few subcellular organanelles in cytoplasm of 2-days-old ΔFvpex13 hyphae (Fig. 7A). Pex14-mCherry is shown as the peroxisomal marker. These results led us to hypothesize that FvPex13 plays a role in peroxisomal protein targeting via its interaction with FvPex5 and FvPex7. We learned that PTS1 receptor FvPex5 predominantly localizes to the cytoplasm, very similar to FvCat, whereas PTS2 receptor FvPex7 and mPTS receptor FvPex19 are mostly found in cytoplasm in ΔFvpex13 mutant (Fig. 7B). These results suggested that FvPex13 function is not only required to maintain localization but also affect the steady-state size and number levels of these docking peroxins affecting peroxisomes.

**Fig. 7.**
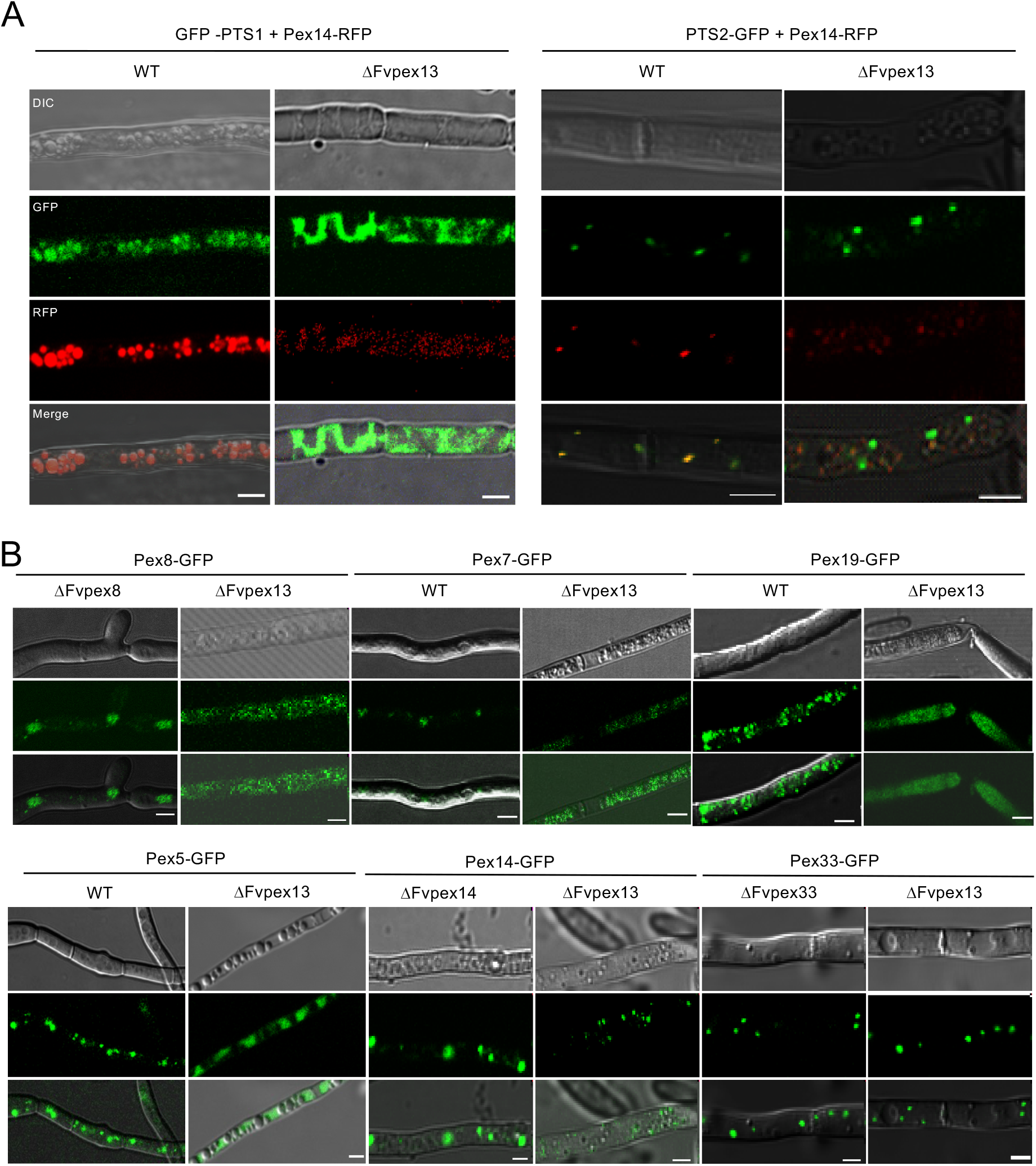
Fvpex13 plays an important role in mycelial peroxin localization. (A) GFP fused to C-terminal PTS1 sequence (PTS1-GFP) and N-terminal PTS2 sequence (GFP-PTS2), localizations were studied in the wild type (WT) and ΔFvpex13 mutant. Pex14-mCherry signal was used as the peroxisomal marker. Bars = 2 μm. (B) Localization of key DTM peroxins Pex8, Pex7, Pex19, and Pex5 were studied in the wild type (WT) and ΔFvpex13 mutant. Pex14 was monitored in ΔFvpex14 and ΔFvpex13, while Pex33-GFP was monitored in ΔFvpex33 and ΔFvpex13.

## Discussion

The key aim of this study was to gain a deeper understanding of how peroxisomal docking/translocation module (DTM) complex impacts peroxin import, the integrity and functionality of peroxisomes, and ultimately fumonisin biosynthesis and virulence in maize pathogen *F. verticillioides*. Interaction between four DTM peroxins, *i.e*. FvPex8, FvPex13, FvPex14, and FvPex33, were demonstrated through Co-IP and yeast two-hybrid assays. Interestingly, FvPex8 exhibited no direct interaction with FvPex14. We also confirmed that FvPex14-mCherry colocalizes with FvPex13-GFP, FvPex8-GFP, and FvPex33-GFP in mycelia, suggesting that these proteins form a complex in *F. verticillioides*. FvPex13 plays a more predominant role in PTS1 than PTS2, though FvPex13 is interactive with both PTS receptors. In *S. cerevisiae*, Pex14 preferentially binds to cargo-loaded Pex5 receptor for pore formation while Pex13 is coupled to the translocation of a cargo protein and the close association of Pex13 and Pex14 is required for peroxisomal matrix protein import (Schell-Steven *et al*., 2005). It was also suggested that Pex13 serves as an initial binding site for the PTS2 pre-import complex, whereas Pex14 assembles with other peroxins in high-molecular-weight complexes after or during dissociation of the PTS2 receptor (Grunau *et al*., 2009, Meinecke *et al*., 2010). Based on literature and our experimental data, we envisaged a schematic model of *F. verticillioides* peroxisomal DTM complex shown in Fig. 8. We suggest that FvPex14 and Fvpex33 prefer the initial recognition of the PTS1 cargo-loaded receptor and PTS2 cargo-loaded receptor respectively, while FvPex13 and FvPex8 prefer to unload cargo-loaded from the receptor. Functional role of Pex8 and its association with other key DTM components has not been characterized in fungal pathogens previously.

**Fig. 8.**
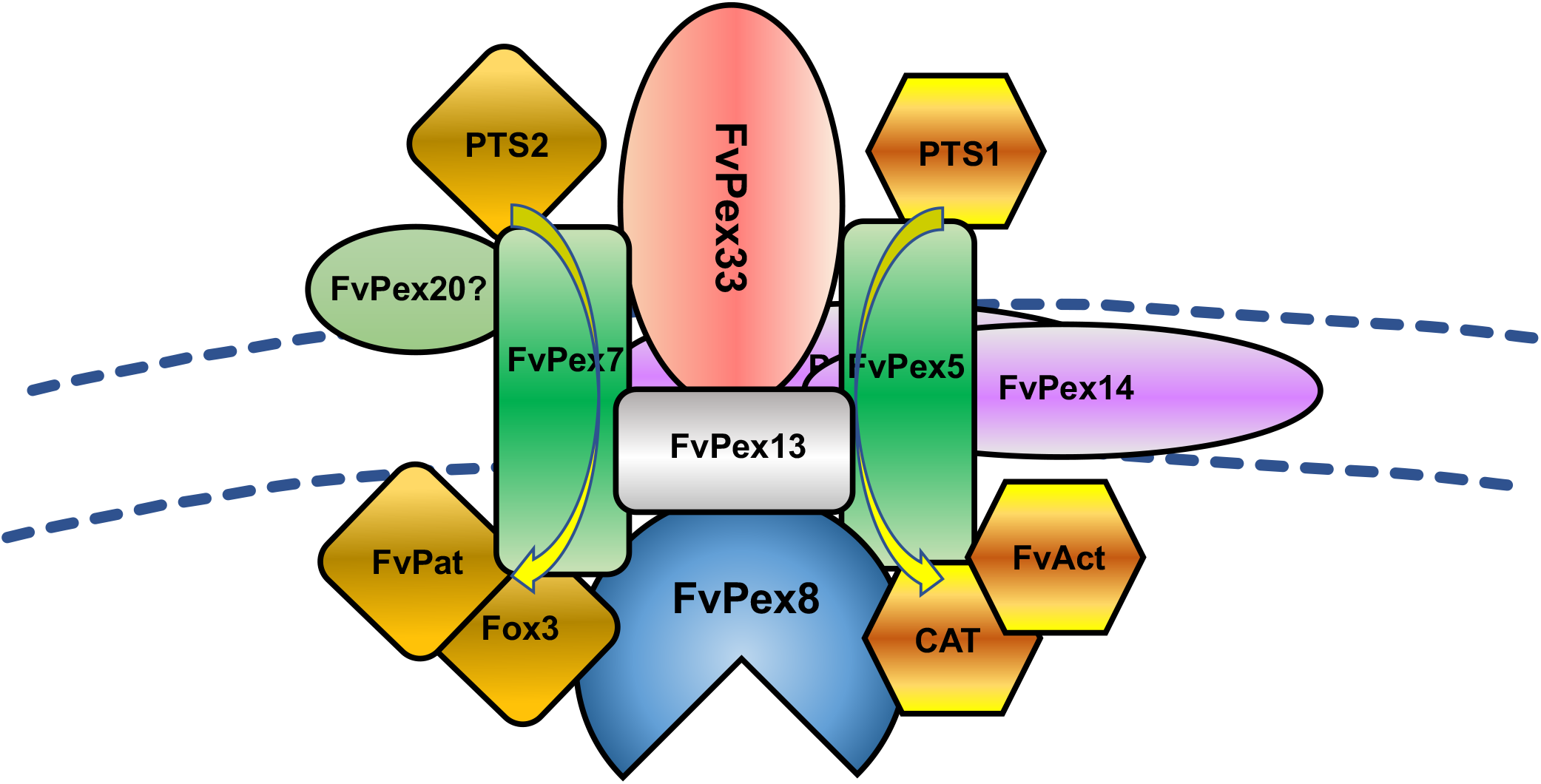
Proposed schematic model of F. ve*rticillioides* peroxisome docking/translocation module (DTM). The proposed model is based on our protein-protein interaction experiments as well as other published fungal models. Our study showed that these proteins are involved in the regulation of peroxisomal docking complex assembly and matrix translocation and ultimately impact various aspects of fungal development, secondary metabolism, and virulence. Notably, we determined that FvPex8 plays an important role in *F. verticillioides* mycotoxin biosynthesis. While we noticed distinct role each component plays, FvPex13 was the key components in regulating virulence, likely in close association with FvPex14 and FvPex33.

Recent studies provide numerous examples how key DTM proteins Pex13 and Pex14 are important not only for peroxisome biogenesis but also for hosts of other cellular and physiological functions in filamentous fungi. Absence of the docking peroxin Pex13 in *Podospora anserina* obstructed sexual development prior to meiosis (Suaste-Olmos *et al*., 2018). *Colletotrichum orbiculare* Pex13 is required for appressorium-mediated plant infection, namely for melanin biosynthesis and turgor generation (Fujihara *et al*., 2010). Chen and colleagues (2018) showed that Pex13 also plays an important role in *Fusarium graminearum* pathogenesis and mycotoxin biosynthesis. We found very similar outcomes in our study, where *F. verticillioides* Pex13 mediated plant infection and severe peroxisomal defects. Pex14, in addition to its key function as a docking protein, plays a main role in formation of two separate import pores, facilitating the PTS1- and PTS2-dependent protein import into the peroxisomal matrix (Montilla-Martinez *et al*., 2015, Meinecke *et al*., 2010). Pex14 has a unique dual function in peroxisome formation and in pexophagy in *Pichia pastoris* and *F. graminearum* (Bellu *et al*., 2001, Farre *et al*., 2008, Deng *et al*., 2013, Lee *et al*., 2017, Chen *et al*., 2018). Pex14 serves an important role in Woronin bodies formation in *Neurospora crassa* and host infection in *F. graminearum* (Managadze *et al*., 2007, Chen *et al*., 2018). In addition to Pex13 and Pex14, another important component of DTM complex in filamentous fungi is Pex33, a chimeric protein consisting of an N-terminal Pex14-like domain and a C-terminal Pex17-like domain. Pex33 has been associated with glyoxysome biogenesis, sexual and asexual development, virulence and secondary metabolism (Peraza-Reyes *et al*., 2011, Li *et al*., 2017, Chen *et al*., 2018, Opalinski *et al*., 2010, Managadze *et al*., 2010). In our study, we found *F. verticillioides* Pex14 and Pex33 collaborate to mediate plant infection and partial peroxisomal defects.

While these key Pex proteins have been extensively investigated in filamentous fungi and other eukaryotes, the role of Pex8 in peroxisomal import machinery and its impact on other physiological roles in pathogenic fungi are still unclear. Pex8, originally discovered as Per1 protein in yeast *Hansenula polymorpha* (syn. *Pichia angusta*), is an intraperoxisomal peroxin that has a PTS1 at the C-terminus and an unusual PTS2 in the mid-protein region (Agne *et al*., 2003, Waterham *et al*., 1994). Pex8 is currently the only recognized intraperoxisomal peroxin, while majority of peroxins are either peroxisomal membrane-bound or in the cytosol (Ma *et al*., 2009, Pieuchot & Jedd, 2012, Girzalsky *et al*., 2010). Pex8 is targeted to peroxisomes by cytosolic receptors Pex5 (PTS1) or Pex7 (PTS2), and the membrane translocation is primarily mediated through docking complex Pex14 (Agne *et al*., 2003, Ma *et al*., 2009). Once positioned in the peroxisomal matrix, Pex8 becomes an important component of importomer complex, anchoring key receptors as well as other membrane-bound peroxins, and serves an important function in both PTS1- and PTS2-dependent protein translocation into peroxisome (Agne *et al*., 2003, Rehling *et al*., 2000). Therefore, it is important to acknowledge that composition and organization of the importomer and docking complex can vary in eukaryotes, not only in species-dependent manner but also in a cargo-dependent manner (Gabaldon, 2010, Heiland & Erdmann, 2005, Lazarow, 2003).

In our study, we learned that FvPex8 plays a crucial role in *F. verticillioides* secondary metabolism but is dispensable for plant pathogenesis. Involvement of peroxisomes in fungal secondary metabolite biosynthesis has been well documented, including in penicillin, AK toxin, paxilline, and deoxynivalenol (DON) (Herr & Fischer, 2014, Saikia & Scott, 2009, Imazaki *et al*., 2010, Martin *et al*., 2012, Chen *et al*., 2018). However, it is still unclear how FvPex8 impacts FB1 biosynthetic pathway or regulation. We learned that the expression of key *FUM* genes are decreased drastically in DFvpex8 mutant, but it is highly unlikely FvPex8 directly regulates genes in the *FUM* cluster. However, it is possible that Pex8, which contains both peroxisomal targeting signals PTS1 and PTS2, functions as an organizer of the peroxisomal protein import machinery. In *S. cerevisiae*, Pex8 is located at the inner surface of the peroxisomal membrane and is essential for the assembly of the importomer (Deckers *et al*., 2010). Acetyl-CoA, one of the key buildings block for the synthesis of polyketides, can be produced via various catabolic pathways, including peroxisomal β-oxidation of fatty acids (Maggio-Hall *et al*., 2005). The FvPex8 mutant showed difficulty growing on multiple carbon sources, raising the hypothesis whether FvPex8 plays a role in peroxisomal carbon recycling and β-oxidation functions. When investigating *F. verticillioides* PTS proteins *in silico*, we also learned that palmitoyl-CoA hydrolase (FVEG_02271, designated FvAct) contains PTS1 and serine palmitoyl transferase (FVEG_12143, designated FvPat) contains PTS2. We can postulate that FvPex8 indirectly impacts hydrolysis of palmitoyl-CoA, which is an acyl-CoA thioester that plays a role in biosynthesis of sphingolipids and in transport of fatty acyl-CoA molecules into the mitochondria for b-oxidation (Brady *et al*., 1969). Based on current DTM models, inner membrane FvPex8 function may impact both import receptors Pex5 (PTS1 cargo shuttle) and Pex7 (PTS2 cargo shuttle) and disrupt import/export of acetyl-CoA necessary for secondary metabolism in *F. verticillioides*. While we observed higher expression levels of *FvPEX5* and *FvPEX7* when we performed retrograde gene regulation of *FvPEX* genes in ΔFvpex8 (Supplementary Information), further study is needed to test this hypothesis.

One unexpected outcome in our study was ΔFvpex8 showing no particular defect in virulence. Studies have shown how key peroxisome biogenesis genes are associated with plant infection and pathogenesis in *Colletotrichum lagenarium, C. orbiculare, M. grisea, A. alternata*, and *F. graminearum* (Ramos-Pamplona & Naqvi, 2006, Wang *et al*., 2007, Imazaki *et al*., 2010, Fujihara *et al*., 2010, Goh *et al*., 2011, Min *et al*., 2012). For instance, CoPex13 is required for appressorium melanization and turgor generation, which are essential for appressorium penetration ability in *C. orbiculare* (Fujihara *et al*., 2010). MoPex33 is required for import of peroxisomal matrix proteins and full virulence in *M. oryzae* (Li *et al*., 2017). A recent publication described how Pex13, Pex14 and Pex33 are critical against host stress agents and pexophagy, ultimately impacting fungal virulence in *F. gramminearum* (Chen *et al*., 2018). Along with these key DTM components, import receptors Pex5 and Pex7 were also shown to be important for fitness and virulence in fungal pathogens (Min *et al*., 2012, Wang *et al*., 2013, Zhang *et al*., 2018a, Goh *et al*., 2011). FvPex8 showed strong physical interaction with FvPex13, therefore it was reasonable to hypothesize that FvPex8 also plays a critical role in pathogenesis. However, this was not the case; only ΔFvpex13 showed impaired virulence in *F. verticillioides* (Fig. 6). FvPex13 is known to affect efficient PTS1 and PTS2 imports, and this requires Pex5-Pex13 interaction *in vivo* (Pires *et al*., 2003, Schell-Steven *et al*., 2005). Loss of peroxisome function may slow pathogen growth due to inefficient utilization of carbon sources from the host and the fungus itself (Fig. 2 and S3). However, we did not observe similar response in host when challenged by ΔPex8 mutant suggesting that Pex8, while it forms a complex with Pex13 and other DTM components, may not directly contribute to virulence in *F. verticillioides*.

To summarize, we identified and characterized *F. verticillioiodes* homologs of DTM components FvPex8, FvPex13, FvPex14, and FvPex33 in this study. Based on our experimental outcomes and literature, we propose a schematic model of core *F. verticillioides* DTM complex (Fig. 8). It is reasonable to hypothesize that these proteins are involved in the regulation of peroxisomal docking complex assembly and matrix translocation and ultimately impact various aspects of fungal development, secondary metabolism, and virulence. However, we clearly noticed a distinct role each component plays. We propose that the DTM complex plays an important role in virulence through FvPex13, likely in association with FvPex14 and FvPex33. For FB1 production, FvPex8 was determined as the key components, but the mechanism in which FvPex8 regulates FB1 biosynthesis needs further investigation.

## Material and methods

### Fungal strains and culture media

*F. verticillioides* wild-type strain 7600 and all transformants (Table S2) were cultured on complete medium (CMII: 2% sucrose, 0.05% magnesium sulfate heptahydrate, 0.1% potassium dihydrogen phosphate, 0.05% potassium chloride, 0.2% nitrate potassium acid, 0.1% hydrolyzed casein, 0.2% peptone, 0.1% Wolfe’s Vitamin Solution [Coolaber, China], 1.8% agar) and minimal medium (MM) (Leslie & Summerell, 2006). CMII agar with 250 μg/ml hygromycin B (Roche, Germany), 200 μg/ml G418 (Sigma, USA) or 800 μg/ml G418 (Sigma) were used for screening transformants. For conidia production assay, six 0.7 cm-diameter agar blocks were added to 3 ml water, vortexed and conidia were quantified by a hemocytometer. For vegetative growth and stress agent sensitivity assays, the wild-type and mutant strains were grown on MM and CMII amended with various carbon sources: potassium acetate (40 mM), sodium butyrate (10 mM), sucrose (2%), palmitic acid (3 mM), oleic acid (3 mM) or linoleic acid (3 mM) (Min *et al*., 2012). Colony diameter was measured after 5 days of incubation. Cell wall integrity was tested by growing the strains on CMII supplemented with 200mg/ml Calcofluor white (CFW) or 0.01% sodium dodecyl sulfate (SDS). The tolerance of strains to exogenous ROS was evaluated by measuring growth on CMII containing 0.05% H_2_O_2_.

### Generation of the DTM gene deletion mutants and GFP/mCherry/Flag-tagged strains

Single DTM gene *F. verticillioides* mutants were generated with a split-marker approach described previously (Zhang *et al*., 2018b). All primers used in this study are listed in Table S3. For double deletion of *FvPEX14* and *FvPEX33*, we replaced the *FvPEX14* with neomycin phosphotransferase (*NEO*) gene in the ΔFvpex33 deletion mutant. Protoplasts were generated, transformed and screened following our standard PCR and qPCR (Yan & Shim, 2020, Zhang *et al*., 2018b). For qPCR, the wild-type and DTM mutants were inoculated in CMII medium for 3 days. The extraction of total RNA and the synthesis of first-strand cDNA were performed as previously described (Yan & Shim, 2020). The expression of peroxins, were detected by qRT-PCR using SuperReal PreMix Plus (SYBR Green) (Tiangen Biotech, China). As an endogenous control, a 200-bp amplicon of *F. verticillioides* β-tubulin (*TUB2*) gene (FVEG_04081) was amplified, and the relative quantification of each transcript was calculated following the standard 2^-ΔΔCT^ method (Yan & Shim, 2020). Primers used to amplify selected genes in qPCR reactions are listed in Table S3. All qPCR assays were conducted in triplicates for each sample and the experiment was repeated three times. Subsequently, we constructed the fusion GFP and mCherry vectors following the protocol described previously (Zhang *et al*., 2018b, Yan & Shim, 2020). All fusion vectors generated were sequenced before transformation. For construction of GFP-tagged strains, we transformed corresponding fusion GFP vectors with G418 sulfate-resistant (*GEN*) gene into protoplasts. For testing whether FvPex13 affected the localization of PTS pathway related genes, we transformed fusion GFP vectors with *GEN* into WT and ΔFvPex13 mutant protoplasts. For testing whether FvPex33 interacts with other DTM, we transformed Flag vectors and fusion Flag-FvPex33 with *GEN* into WT and ΔFvPex33 mutant protoplasts, respectively.

### ROS and lipid staining and quantification

For lipid staining, 2-day-old mycelia were harvested as 0h, and corresponding 2-day-old mycelia were transfered to PBS medium for 18 h, washing mycelia twice with water. Hypha was tested by staining with Nile Red on 0h and 18 h respectivly (Nguyen *et al*., 2011). Cellular ROS accumulation in the fungal hyphae was analyzed using the CM-H2DCFDA staining. Mycelia were harvested and washed with PBS for 3 times. Hypha was stained with CM-H2DCFDA (5 μM) (Sigma) for 10 min in the dark following GENMED reagent protocols, scanning confocal microscope with excitation at 490nm (GENMED Scientifics, USA). Accumulation of ROS in mycelia was visualized with 6-chloromethyl-2’,7’-dichlorodihydrofluorescein diacetate (CM-H2DCFDA) staining. Endogenous ROS scavenging capacity was tested by growing the strains on CMII medium 5 days, and the colony were stained with 3,3’-diaminobenzidine (DAB) for H_2_O_2_ detection and with 2mM Nitro Blue Tetrazolium (NBT) (Sigma) in 20mM phosphate buffer, pH6 for O^2-^ detection treated, respectively.

### Co-immunoprecipitation and yeast two-hybrid assays

Strains were grown in CMII medium for 2 days before immunoblot and immunoprecipitation analyses were conducted as previously described (Zheng *et al*., 2015). For total protein extraction, mycelia of FvPex13-GFP and FvPex14-mCherry were ground into a fine powder in liquid nitrogen and resuspended in 30-mL extraction buffer (10 mM Tris/Cl pH 7.5, 150 mM NaCl, 0.5 mM EDTA, 0.5% NP40, 1 mM PMSF). The supernatants were centrifuged 15,000 g for 25 min at 4°C to remove cell debris. For immunoprecipitation of GFP-fusion proteins from cellular extracts, equal concentration of total proteins was isolated and incubated with 20~30 μL of GFP-Trap agarose beads (ChromoTek, Germany) according to manufacturer’s instructions. Proteins eluted from GFP-Trap agarose beads were analyzed by immunoblot detection with the anti-mCherry, anti-Flag (Clontech, USA) and anti-GFP (Sigma) antibodies.

The yeast two-hybrid assay was carried out following our previous standard protocol (Zhang *et al*., 2018b). The coding sequence of each tested gene was amplified from wild-type cDNA. *FvPEX13* and *FvPEX14* were cloned into pGBKT7 as the bait vectors. Other components of the DTM *FvPEX8, FvPEX14, FvPEX33*, along with *FvPEX5* and *FvPEX7* cDNA were amplified and cloned into pGADT7 as the prey vectors. The pairs of plasmids pGBKT7-P53 and pGADT7-T, pGBKT7-Lam and pGADT7-T were used as the positive and the negative control, respectively. All primers used in this study were listed in Table S3. Further details are provided in Supplementary Information.

### Virulence and FB1 assays

Infection assays on maize and sugarcane stalks were conducted as previously described (Zhang *et al*., 2018b, Zhang *et al*., 2015). Subsequently, maize and sugar cane stalks were split longitudinally to assess disease severity and to quantify the extent of the rot by Image J software. For FB1 assay, conidia were inoculated on surface sterilized B73 corn kernel for 10 days and quantified with Fumonisin B1 ELISA Kit following manufacturer’s suggested protocol (Biovision, USA). FB1 concentrations were obtained by the optical density (OD value) of each well at 450 nm with a microplate reader by comparing to FB1 standard curve. *F. verticillioides* biomass quantification was calculated by qPCR based on the *TUB*2 standard curve, which was measured at 0.01, 0.1, 1.0, 10 ng/μl concentrations (Rocha *et al*., 2015). Sample FB1 levels were normalized by calculating FB1 ppm/ *TUB*2 DNA concentration. To compare the *FUM* gene expression, RNA samples were prepared from ΔFvpex8 mutant and *F. verticillioides* strains grown on corn kernel for 10 days. The expression levels of *FUM1, FUM8, FUM19* and *FUM21* were normalized to *TUB2* expression. The experiment was repeated three times.

## Author Contributions

W.Y., W-B.S. and Z.W. conceived and designed the experiments. W.Y., M.L., H.Y. and Y.H. performed the experiments. W.Y., J.Z., G.L., W-B.S. and Z.W analyzed the data. W.Y., H.Y., W-B.S. and Z.W. wrote the paper.

## Acknowledgments

This work was supported by the NSFC grants (#31601599) and Science and Technology Innovation Special Fund of FAFU(#CXZX2017138). The authors declare no conflict of interest.

## SUPPLEMENTARY TABLES

**Table S1.** Putative *F. verticillioides* DTM components identified by Blastp analysis

**Table S2.** *F. verticillioides* strains used in this study

**Table S3.** PCR primers used in this study

## SUPPLEMENTARY FIGURES

**Figure S1. Sequence analysis of Docking peroxin homologs in *F. verticillioides***

(A) Schematic description of conserveds domain in peroxins FvPex13, FvPex14 and FvPex33. (B) Phylogenetic relationship of FvPex33 and its homologs calculated with neighbor joining method using the MEGA 6.0 program according to the alignment. (C) Possible transmembrane helices predicted in FvPex8, FvPex13, FvPex14, and FvPex33 analyzed by TMPRED algorithm.

**Figure S2. Docking protein interaction by yeast two hybrid assays.**

Interactions between DTM peroxins were tested using the yeast two-hybrid assay. Yeast transformants expressing plasmids harboring the indicated constructs were assayed for growth on selective plates (SD/Leu/Trp/His/Ade), and for b-galactosidase (LacZ) activities. The pGBKT7-53/pGADT7-T and pGBKT7-Lam/pGADT7-T was used as positive and negative control, respectively.

**Figure S3. Vegetative growth and growth inhibition of mutants on CMII agar.**

(A) The colony morphology of DTM mutants and the wild-type strain on CMII agar and CMII amended with linoleic acid. (B) Mycelial growth inhibition rates were calculated by [(the diameter of WT strain - the diameter of mutant strain)/(the diameter of WT strain)]x100%. (C) conidiation in the wild type and DTM mutants on CMII agar. An asterisk above the column indicates a statistically significant difference (P<0.05) as anlyzed by t-test.

**Figure S4. Reduction in lipid droplet degradation and carbon sources utilization**

(A) Wild-type (WT) and mutant strains were grown for 2 days in rich medium, then transferred to 1% potassium acetate medium and stained with Nile red to observe the lipid droplet formation of the hyphae after starvation at 0 h and 18 h time points. (B) Lipid droplet gross area was measured in each sample at 0 h and 18 h time points. (C) Lipid droplet sizes were measured measured in each sample at 0 h and 18 h time points. Bars denote standard deviations from three repeated experiments. An asterisk above the column indicates a statistically significant difference (P<0.05) as anlyzed by t-test.

**Figure S5. Role of DTM peroxins in sugarcane stalk rot virulence.**

(A) Sugarcane stalks (cultivar Badila) were inoculated with fungal spore suspension (10^6^/ml) at the internodal region and incubated in room temperature for 7 days. Subsequently, stalks were split longitudinally to visually inspect rot symptoms. (B) Image J software was used to quantify the extent of the rot in each sample. Three independent biological repetitions were performed and error bars represent the standard deviation from three replicates. An asterisk above the column indicates a statistically significant difference (P<0.05) as anlyzed by t-test. (C) Sterile water was used as a negative control on maize stalk.

## SUPPLEMENTARY INFORMATION

Supplementary Results and Supplementary Methods are provided in a separate document

## Notes

### Competing Interest Statement

The authors have declared no competing interest.

### Summary of Updates

The manuscript has been revised to place more emphasis on the role of FvPex8 in fumonisin biosynthesis in maize pathogen Fusarium verticillioides. While paying a close attention to two core components of the docking complex, Pex13 and Pex14, we expanded our study into additional peroxisomal proteins, namely Pex8, that have not been well characterized in plant pathogenic fungi

