## Supplementary Figures for "*Fusarium verticillioides* FvPex8 is a key component of peroxisomal docking/translocation module that serves important roles in fumonisin biosynthesis but not in virulence"

Yu et al. Fig. S1

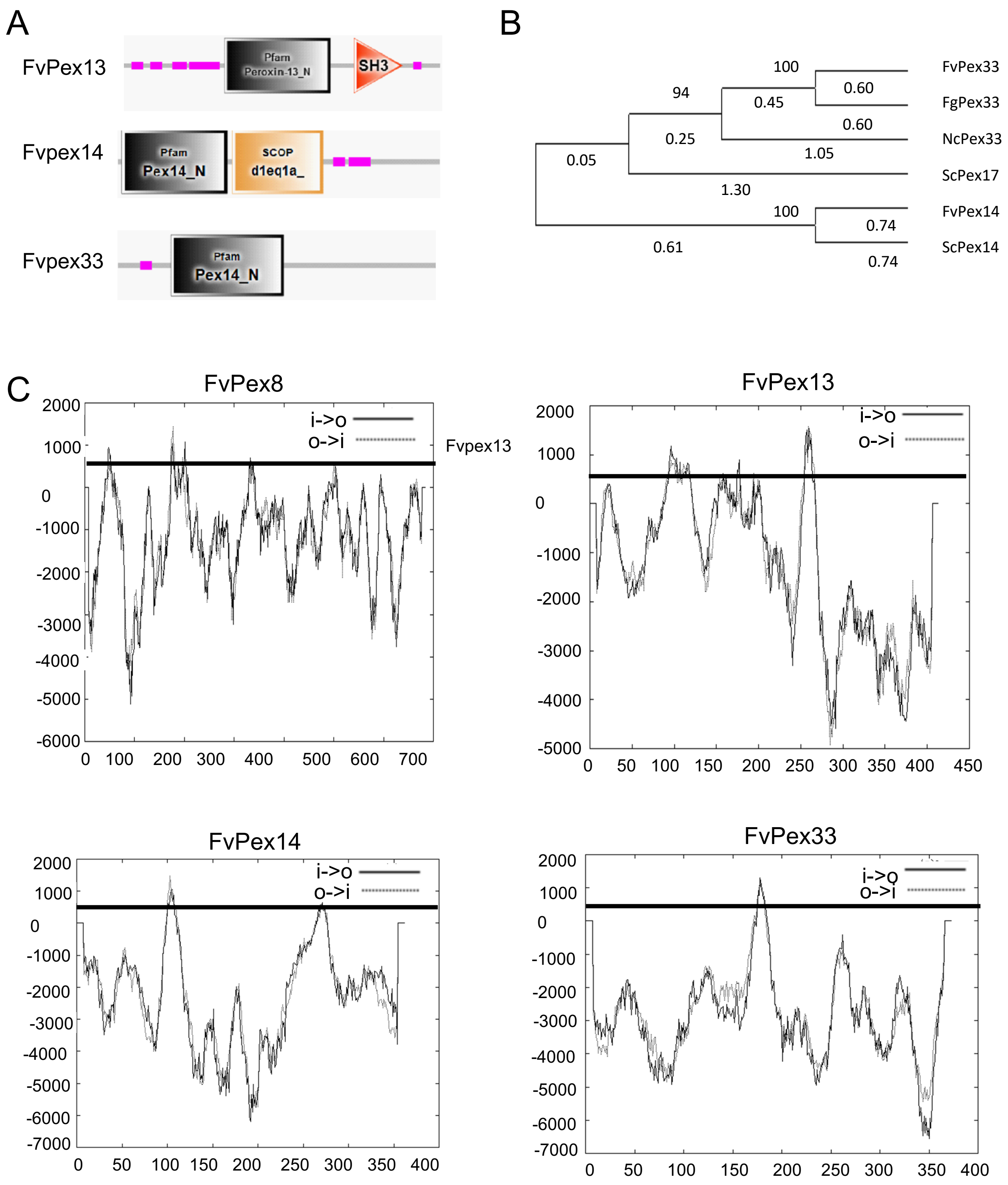


http://www.ch.embnet.org/software/TMPRED_form.html Possible transmembrane helices :Only scores above 500 are considered significant.

Yu et al. Fig. S2

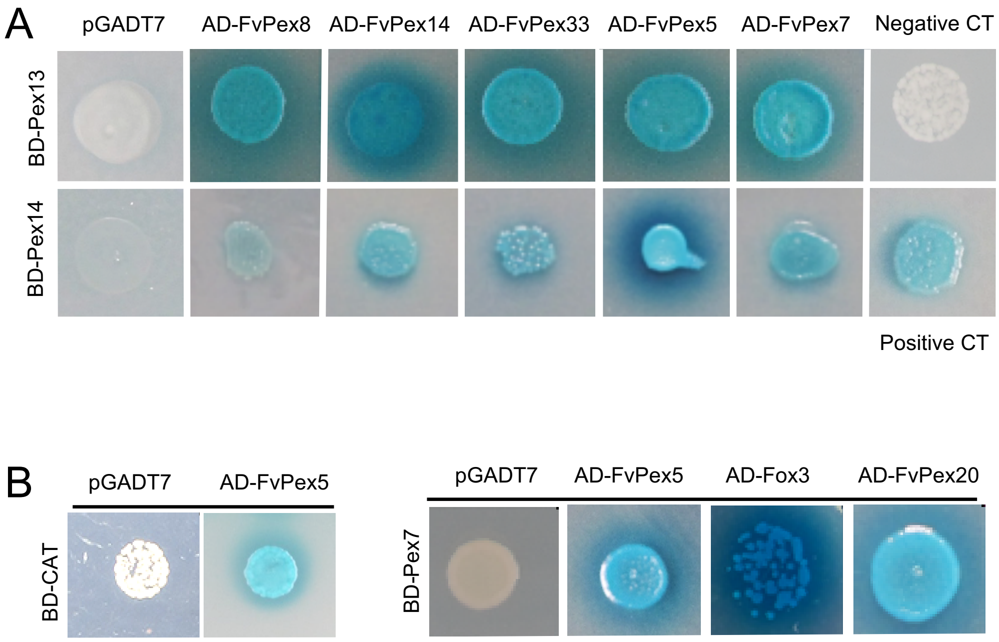

Yu et al. Fig. S3

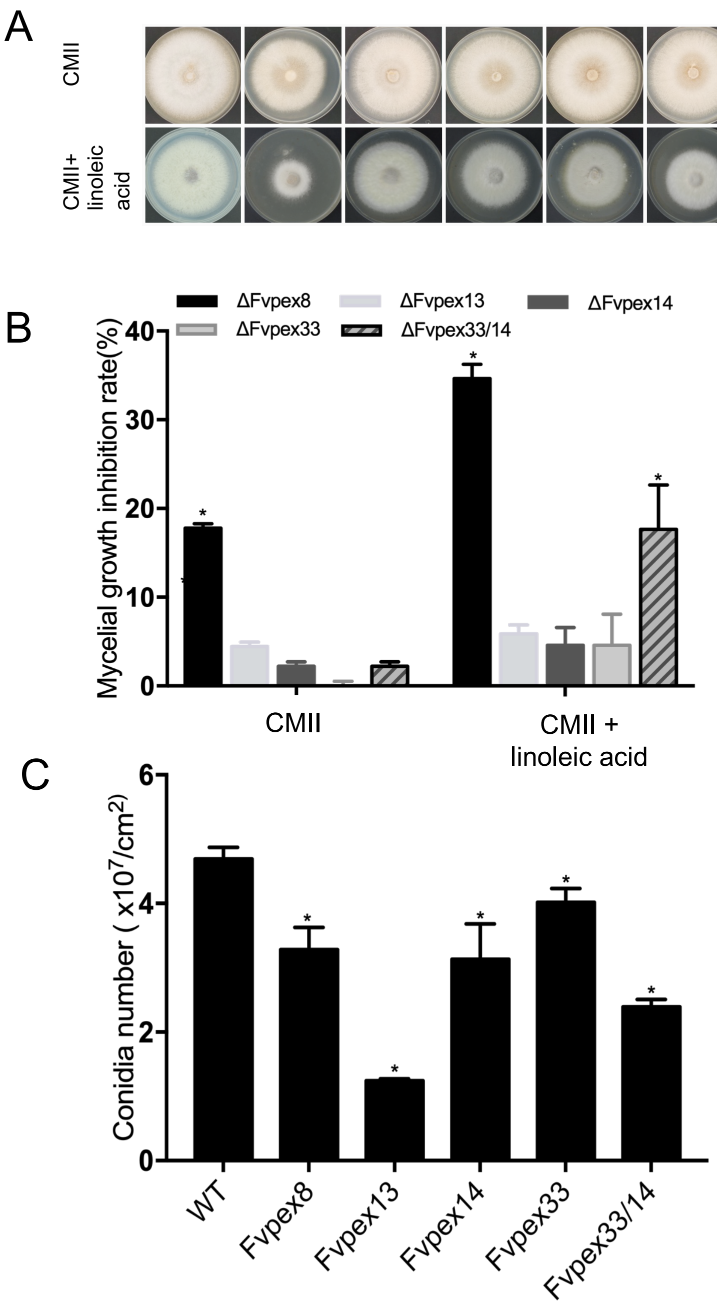

Yu et al. Fig. S4

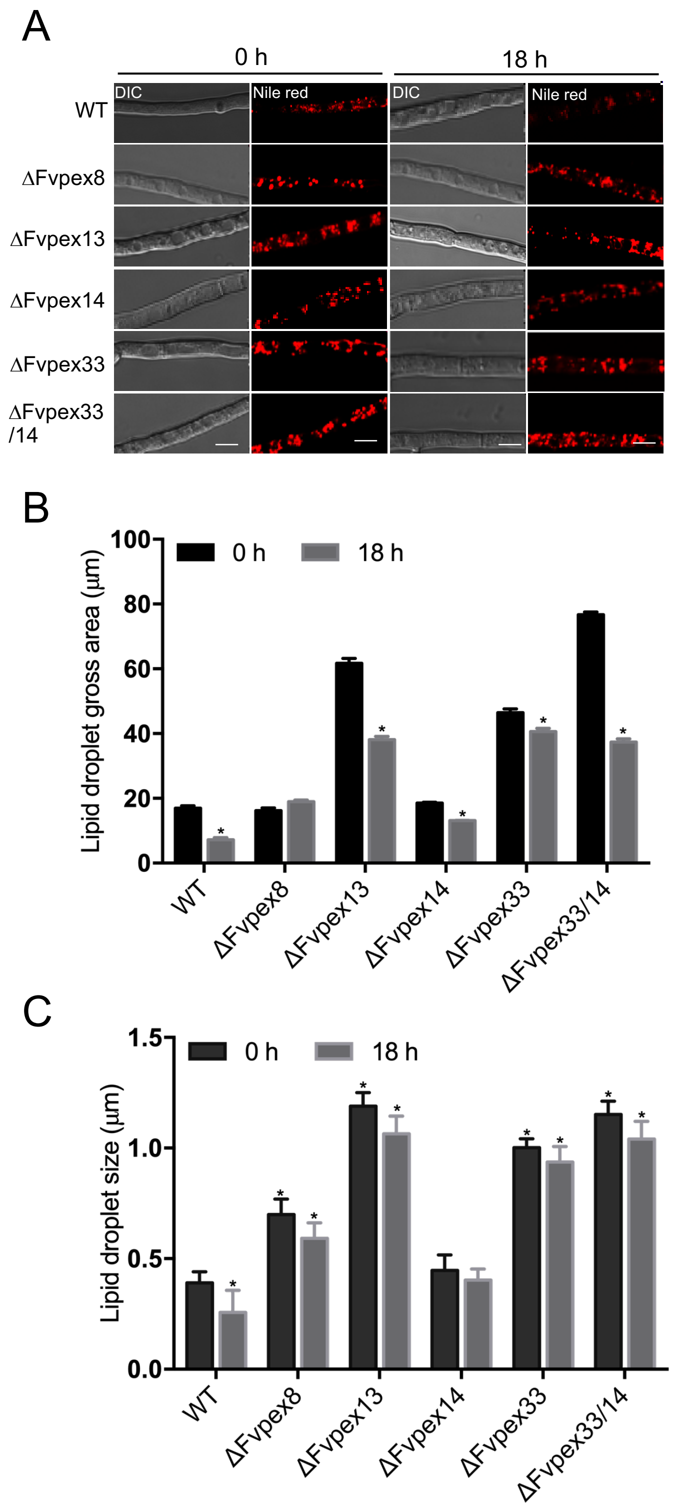

Yu et al. Fig. S5

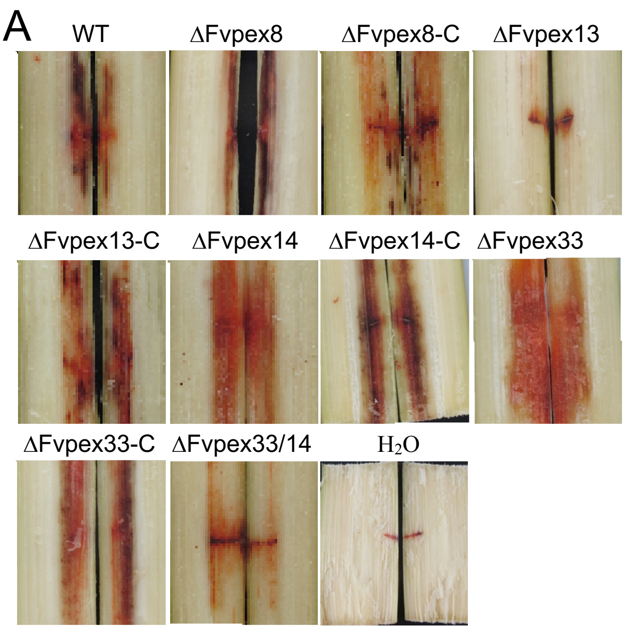


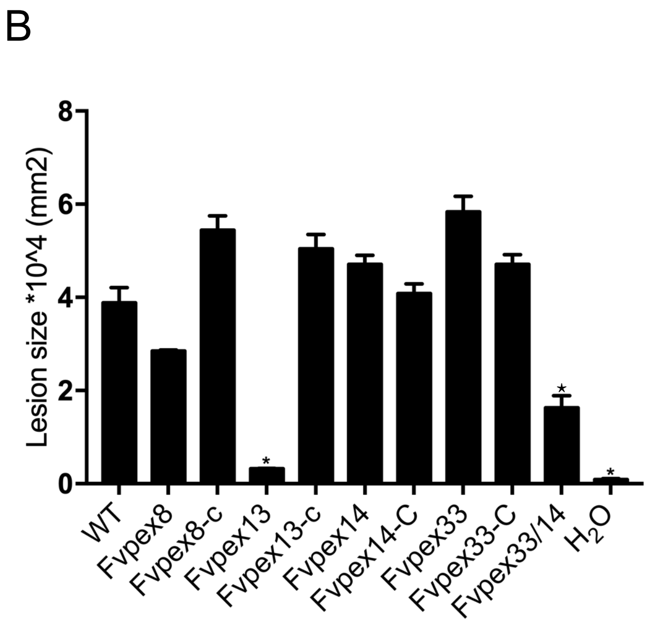


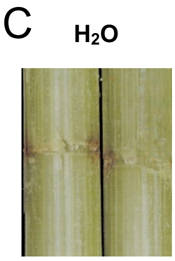
