## Supplementary Information for "*Fusarium verticillioides* FvPex8 is a key component of peroxisomal docking/translocation module that serves important roles in fumonisin biosynthesis but not in virulence"

**Supplementary Results**

**Results S1. Sequence analyses of *F. verticillioides* DTM peroxins**

We identified FVEG_00529 (designated FvPex13) with two functional motifs Peroxin-13_N and SH3 (Fig. S1 A), which showed 44% sequence identity to the *Saccharomyces cerevisiae* Pex13 (Table S1). The locus FVEG_03847 (designated FvPex14), contained two functional motifs Pex14_N and SCOP (Fig. S1 B) and showed 32% sequence identity to the *S. cerevisiae* Pex14 (Table S1). The locus FVEG_05423 (designated FvPex8), with SKL (PTS1) at C-terminus and RL-X5-QH (PTS2) near the N-terminus, showed 47% sequence identity to the *N. crassa* Pex8 (Table S1). Lastly, the locus FVEG_11334 (designated FvPex33) was predicted to harbor one functional motif Pex14_N at the N terminus (Fig. S1 C) and showed 32% sequence identity to the *N. crassa* Pex33 (Table S1). Further comparison of FvPex33 against Pex14- and Pex17-like proteins in *S. cerevisiae*, *F. graminearum*, *F. verticillioides* strongly suggested that FvPex33 is distinct from Pex14 and Pex17 and is a unique component of DTM in *F. verticillioides* (Fig. S1 D).

**Results S2. Retrograde gene regulation in Peroxins**

We first examined whether expression of other peroxin genes were affected in the DTM mutants (Method S4). These included four DTM genes (*FvPEX8*, *FvPEX13*, *FvPEX14* and *FvPEX33*), two PTS receptor genes (*FvPEX5* and *FvPEX7*), and a membrane protein receptor gene (*FvPEX19*). *FvPEX8* exhibited no detectable change when tested in three other DTM mutants. *FvPEX13* showed decreased expression while *FvPEX14* and *FvPEX33* showed drastically increased expression in ΔFvpex8 mutant. *FvPEX5* and *FvPEX7* genes showed elevated transcription levels in ΔFvpex8 mutant while showing decreased transcription in ΔFvpex33/14 mutant when compared with the wild type. Meanwhile, *FvPEX19* was suppressed in ΔFvpex8, ΔFvpex33, ΔFvpex33/14 mutants. When comparing DTM mutants, deletion of *FvPEX8* led to most noteworthy changes in peroxin gene expression, particularly dramatic overexpression of PTS receptor genes (*FvPEX5* and *FvPEX7*) as well as two DTM genes (*FvPEX14* and *FvPEX33*) led us to hypothesize that FvPex8 plays an important role in maintaining peroxisomal membrane assembly and cargo transport machinery. It is also notable that ΔFvpex33/14 double mutant, but not ΔFvpex14 or ΔFvpex33 single mutant, exhibited strong negative impact on expression of *FvPEX5,* *FvPEX7,* *FvPEX14* and *FvPEX33.*

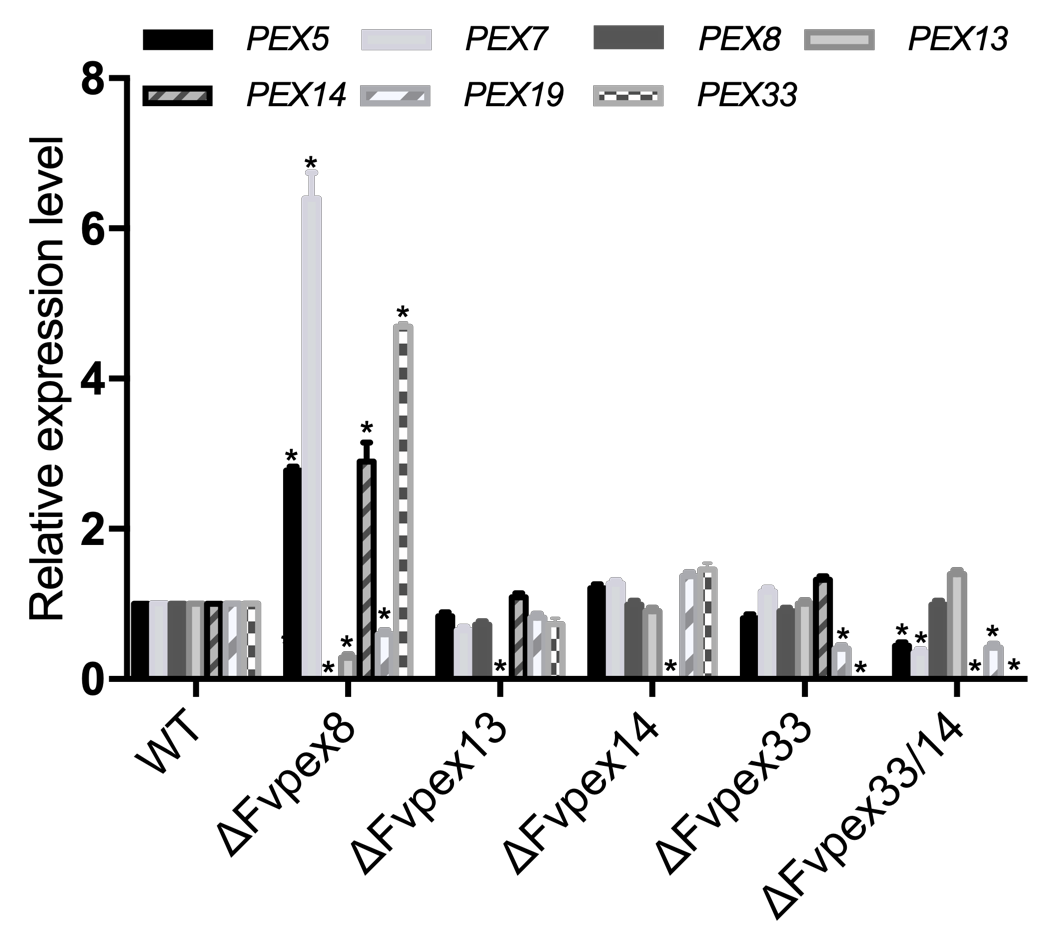

Expression changes of Peroxin genes were studied in each mutant using qPCR. Relative transcript abundance was measured by qPCR and normalized to *F. verticillioides* β-tubulin gene. Values are means ± SE of three replicates. An asterisk above the column indicates a statistically significant difference (P<0.05) as analyzed by t-test.

**Results S3. Lipid droplet degradation in peroxisome DTM complex mutants**

In an earlier study, Deng *et al* (2016) showed that a mutation in peroxin gene *MoPEX1* can delay lipid mobilization and degradation during appressorium development *in M. oryzae*. The lipid degradation defects were also observed in conidia of *F. graminearum* DTM mutants (Chen *et al*, 2018). To examine whether *F. verticillioides* peroxins are involved in lipid degradation, we used Nile Red to stain intracellular lipid droplets. We observed how WT and DTM mutants accumulated lipid droplets when cultured in rich CMII medium. The mutants accumulated higher levels of lipid droplets compared to WT, especially in ΔFvpex13 and ΔFvpex14/33 and (Fig. S4 A). When strains were shifted to starvation conditions for 18 h, the average size of lipid droplets in WT cells was significantly reduced but not in the mutants (Fig. S4 B and S4 C). The relative lipid droplet gross area was significantly reduced in ΔFvpex33 and ΔFvpex13, but these two mutants showed much higher gross area at 0 h when compared to WT and other mutants (Fig. S4 B). These data suggested that DTM complex is important for lipid turnover under starvation conditions.

### Deng, S., Gu, Z., Yang, N., Li, L., Yue, X., Que, Y., Sun, G., Wang, Z., and Wang, J. (2016). Identification and characterization of the peroxin 1 gene MoPEX1 required for infection-related morphogenesis and pathogenicity in *Magnaporthe oryzae*. *Sci. Rep*. 8 (6):36292.

**Chen, Y., Zheng, S., Ju, Z., Zhang, C., Tang, G., Wang, J., Wen, Z., Chen, W. and Ma, Z.** (2018) Contribution of peroxisomal docking machinery to mycotoxin biosynthesis, pathogenicity and pexophagy in the plant pathogenic fungus *Fusarium graminearum*. *Environmental Microbiology,* **20,** 3224-3245.

**Supplementary Methods**

**Method S1. Targeted gene replacement strategy**

Each targeted gene was successfully replaced with hygromycin resistance cassette (*HYG*) using the split marker technique in *F. verticillioides* wild type strain (Zhang et al. 2018). ΔFvpex14/33 mutant was generated through targeted gene replacement with the geneticin resistance cassette in the ΔFvpex33 background.

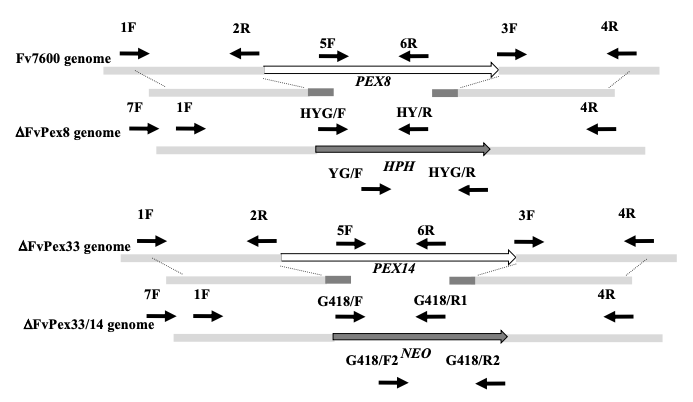

After transformation, hygromycin-resistant transformants were first screened by PCR with designated primer pairs 5F/6R and 11F/8R to confirm the gene deletion and further characterized by Southern blot with the Digoxigenin High Prime DNA Labeling and Detection Starter Kit I (Roche). Southern hybridization results further confirmed that the constructs replaced gene in the genome. The anticipated band sizes in Southern hybridization were; 1) Using *FvPEX13* 3’-flanking region as the probe and digested with *BamH*I, 5.0-kb band in ΔFvpex13 and 6.8-kb band in WT; 2) Using *FvPEX14* 3’-flanking region as the probe and digested with *EcoR*I, 5.4-kb band in ΔFvpex14 and ΔFvpex14/33 and 2.8-kb band in WT; 3) Using FvPEX8 3’-flanking region as the probe and digested with *Sal*I, 3.9 kb band in ΔFvpex8 and 0.9-kb band in WT. In the ΔFvpex14/33 double mutant, the marker gene (*NEO*) was amplified from the plasmid pKNTG (Zheng et al. 2012) with primer pairs of G418-F1/ G418-R1 and G418-F2/ G418-R2, respectively (Table S2). After transformation, G418 sulfate-resistant transformants were first screened by PCR with designated primer pairs 5F/6R and 11F/8R to confirm the *Fvpex14* gene deletion and further verified by qPCR.

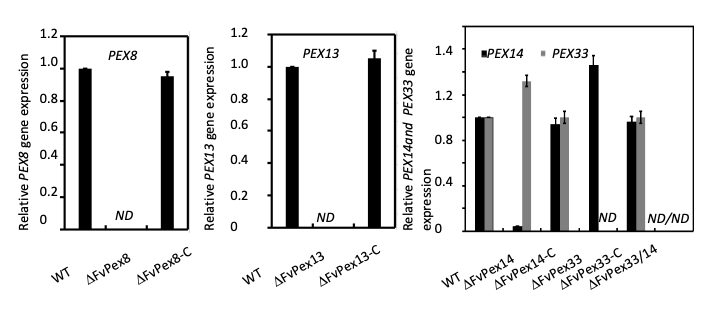

**Method S2. GFP and mCherry tagged strains**

For *in vivo* localization study, we generated GFP strains by introducing FvPex5::GFP, FvPex7::GFP, FvPex8::GFP, FvPex14::GFP, FvPex19::GFP, FvPex33::GFP, and FvPex14::mCherry fusion constructs with their endogenous promoter. For FvPex13::GFP, we used a construct with RP27 promoter, since the construct with the native promoter failed to provide good fluorescence signal. For *in vivo* co-localization, wild-type protoplasts were transiently transformed with two plasmids coding for FvPex13-GFP and FvPex14-mCherry. We constructed other strains with FvPex13::GFP and FvPex14::mCherry, FvPex33::GFP and FvPex14::mCherry, FvPex8::GFP and FvPex14::mCherry in wild-type protoplasts. GFP::FvCat, FvFox3::GFP and mCherry::FvPex14 were also constructed and introduced into *F. verticillioides* protoplasts.

**Method S3. Yeast two-hybrid assay**

The yeast two-hybrid assay was carried out using Matchmaker GAL4 Two-Hybrid System 3 (Clontech) according to the manufacturer’s instructions. The coding sequence of each tested gene was amplified from the cDNA of Fv7600. Here, *FvPEX13*, *FvPEX14,* and *FvPEX33*, and *FvPEX8* were cloned into pGBKT7 as the bait vectors pGBKT7-FvPEX13, pGBKT7-FvPex14, pGBKT7-FvPex33, and pGBKT7-FvPex8, respectively. Other components of the DTM *FvPEX33*, *FvPEX14*, *FvPEX8*, *FvPEX5*, *FvPEX7* cDNA were amplified and cloned into pGADT7 as the prey vectors. The resulting yeast two hybrid plasmids, which were verified by sequencing, were co-transformed into the yeast strain AH109 (Clontech) according to LiAc/SS-DNA/PEG transformation procedure. The pairs of plasmids pGBKT7-P53 and pGADT7-T, pGBKT7-Lam and pGADT7-T were used as the positive control and the negative control, respectively. The Leu+ and Trp+ yeast transformants were isolated and assayed for growth on SD-Trp-Leu-His-Ade medium with X-α-gal at specified concentrations. All primers used in this study were listed in supporting information Table S3.

**Zhang, H., Mukherjee, M., Kim, J. E., Yu, W. and Shim, W. B.** (2018) Fsr1, a striatin homologue, forms an endomembrane-associated complex that regulates virulence in the maize pathogen *Fusarium verticillioides*. *Mol Plant Pathol,* **19,** 812-826.

**Zheng, W., Zhao, X., Xie, Q., Huang, Q., Zhang, C., Zhai, H., Xu, L., Lu, G., Shim, W.-B., and Wang, Z.** (2012) A Conserved Homeobox Transcription Factor Htf1 Is Required for Phialide Development and Conidiogenesis in *Fusarium* Species. *PLoS One* **7**, e45432.

**Method S4. Expression analyses of peroxin genes**

For qRT-PCR, the wild-type and DTM mutants were inoculated in CMII medium for 3 days. The extraction of total RNA and the synthesis of first-strand cDNA were performed as previously described (Yan and Shim, 2020). The expression of peroxins, were detected by qRT-PCR using SuperReal PreMix Plus (SYBR Green) (Tiangen Biotech, China). As an endogenous control, a 200-bp amplicon of *F. verticillioides* β*-*tubulin (*TUB2*) gene (FVEG_04081) was amplified, and the relative quantification of each transcript was calculated following the standard 2^-ΔΔCT^ method (Yan and Shim, 2020). Primers used to amplify selected genes in qPCR reactions are listed in Table S3. All qPCR assays were conducted in triplicates for each sample and the experiment was repeated three times.
