## Supplementary Table 1 for "*Fusarium verticillioides* FvPex8 is a key component of peroxisomal docking/translocation module that serves important roles in fumonisin biosynthesis but not in virulence"

**Table S1. Putative DTM components were identified by Blastp analysis**

| Query Protein | Homologous Locus in *F. verticillioides* | E-value | Sequence Identity |
|---|---|---|---|
| PEX13 [Saccharomyces cerevisiae] KZV09435.1 | FVEG_00529 (FvPex13) | 3e-69 | 44% |
| PEX14 [Saccharomyces cerevisiae] KZV11075.1 | FVEG_03847 (FvPex14) | 2e-40 | 32% |
| PEX33 [*Neurospora crassa OR74A*] XP_956984.3 | FVEG_11334 (FvPex33) | 3e-64 | 32% |
| PEX8 [*Neurospora crassa OR74A*] XP_957019.3 | FVEG_05423 (FvPex8) | 0 | 47% |
| Pex5 [ *Neurospora crassa OR74A*] XP_965347.3 | FVEG_01301(FvPex5) | 0 | 63% |
| Pex7 [*Saccharomyces cerevisiae*] EDV08167.1 | FVEG_09607 (FvPex7) | 3e-18 | 30% |
| Pex19 [*Neurospora crassa OR74A*] XP_961091.1 | FVEG_02652 (FvPex19) | 2e-88 | 55% |
| Carnitine O-acetyltransferase [*Neurospora crassa OR74A*] XP_962672.2 | FVEG_07327 (FvCAT) | 3e-74 | 34% |
| 3-ketoacyl-CoA thiolase [ *Neurospora crassa OR74A*] XP_958712.1 | FVEG_10433 (FvFOX3) | 0 | 81% |
