## Supplementary Table 2 for "*Fusarium verticillioides* FvPex8 is a key component of peroxisomal docking/translocation module that serves important roles in fumonisin biosynthesis but not in virulence"

**Table S2. Wild-type and mutant strains of fungi used in this study**

| Strain | Genotype description | Reference |
|---|---|---|
| Δ*Fvpex13* | FVEG_00529 deletion mutant of Fv7600 | This study |
| Δ*Fvpex13-Com* | *ΔFvpex13* strain expressing the Fvpex13-GFP construct | This study |
| Δ*Fvpex14* | FVEG_03847 deletion mutant of Fv7600 | This study |
| Δ*Fvpex14-Com* | *ΔFvpex14* strain expressing the Fvpex14-GFP construct | This study |
| Δ*Fvpex33* | FVEG_11334 deletion mutant of Fv7600 | Zhang et al 2018 |
| Δ*Fvpex33-Com* | *ΔFvpex33* strain expressing the Fvpex33-GFP construct | This study |
| ΔΔ*Fvpex33/14* | FVEG_03847 deletion mutant in Δ*Fvpex33* background | This study |
| Δ*Fvpex8* | FVEG_5423 deletion mutant of Fv7600 | This study |
| Δ*Fvpex8-Com* | *ΔFvpex8*strain expressing the Fvpex8-GFP construct | This study |
| Δ*Fvpex13/*Pex13:GFP:pPex14:RFP | *ΔFvpex13* strain expressing the Fvpex13-GFP *and* Pex14:mcherry construct | This study |
| Δ*Fvpex8/*Pex8:GFP:pPex14:RFP | *ΔFvpex8* strain expressing the Fvpex13-GFP and Pex14:mcherry construct | This study |
| Δ*Fvpex33/*Pex33:GFP:pPex14:RFP | *ΔFvpex33* strain expressing the Fvpex33-GFP and Pex14:mcherry construct | This study |
| Δ*Fvpex33/*Pex33:GFP:pPex13:RFP | *ΔFvpex8*strain expressing the Fvpex33-GFP and Pex13:mcherry construct | This study |
| WT*/*Pex13:GFP:pPex5:RFP | Fv7600 strain expressing the Fvpex13-GFP nd Pex5:mcherry construct | This study |
| WT/Pex14:GFP:pPex5:RFP | Fv7600strain expressing the Fvpex14-GFP and Pex5:mcherry construct | This study |
| WT*/*Pex33:GFP:pPex5:RFP | Fv7600strain expressing the *Fvpex33*-GFP and Pex5mcherry construct | This study |
| WT*/*Pex8:GFP:pPex5:RFP | Fv7600 expressing the Fvpex8-GFP and Pex5:mcherry construct | This study |
| Δ*Fvpex13/*Pex14:GFP | *ΔFvpex13* strain expressing the Fvpex14 GFP construct | This study |
| Δ*Fvpex13/*Pex33:GFP | *ΔFvpex13* strain expressing the Fvpex33 GFP construct | This study |
| Δ*Fvpex13/*Pex8:GFP | *ΔFvpex13* strain expressing the Fvpex8 GFP construct | This study |
| Δ*Fvpex13/*Pex19:GFP | *ΔFvpex13* strain expressing the Fvpex19 GFP construct | This study |
| Δ*Fvpex13/*Pex5:GFP | *ΔFvpex13* strain expressing the Fvpex5 GFP construct | This study |
| Δ*Fvpex13/*Pex7:GFP | *ΔFvpex13* strain expressing the Fvpex7 GFP construct | This study |
| Δ*Fvpex13/*FOX3:GFP: Pex14:RFP | *ΔFvpex13* strain expressing the FvFOX3:GF-GFP and Pex14:mcherry construct | This study |

| | | |
|---|---|---|
| Δ*Fvpex13*/CAT:GFP: Pex14:RFP | *ΔFvpex13* strain expressing the Fv CAT -GFP and Pex14:mcherry construct | This study |
| Δ*Fvpex13*/Pex19:GFP: Pex14:RFP | *ΔFvpex13* strain expressing the Fvpex19-GFP and Pex14:mcherry construct | This study |
| WT /Pex14:GFP | Fv7600strain expressing the Fvpex14 GFP construct | This study |
| WT /Pex33:GFP | Fv7600 strain expressing the Fvpex33 GFP construct | This study |
| WT/Pex8:GFP | Fv7600 strain expressing the Fvpex8 GFP construct | This study |
| WT/Pex19:GFP | Fv7600 strain expressing the Fvpex19 GFP construct | This study |
| WT/Pex5:GFP | Fv7600 strain expressing the Fvpex5 GFP construct | This study |
| WT/Pex7:GFP | Fv7600 strain expressing the Fvpex7 GFP construct | This study |
| WT/FOX3:GFP: Pex14:RFP | Fv7600 strain expressing the FvFOX3:GF-GFP and Pex14:mcherry construct | This study |
| WT/CAT:GFP: Pex14:RFP | Fv7600 strain expressing the Fv CAT -GFP and Pex14:mcherry construct | This study |
| WT/Pex19:GFP: Pex14:RFP | Fv7600 strain expressing the Fvpex19-GFP and Pex14:mcherry construct | This study |
| WT/ Pex14:RFP | Fv7600 strain expressing Pex14-mcherry | This study |
| WT/ Pex33:RFP | Fv7600 strain expressing Pex33-mcherry | This study |
