## Supplementary Table 3 for "*Fusarium verticillioides* FvPex8 is a key component of peroxisomal docking/translocation module that serves important roles in fumonisin biosynthesis but not in virulence"

**Table S3. PCR primers used in this study.**

| Primers | Sequence (5'-3') | Application |
|---|---|---|
| Pex13 AFF | CGAGTCCCAAATGAGAAGTG | ΔFvpex13 deletion and probe |
| Pex13 AFR | ATCCTGCCAGCCTACCCTAT | |
| Pex13 BFF | GGCTATCTTTGCCCAGTTTC | |
| Pex13 BFR | ATCGTTATTGCCCTCACCTT | |
| Pex13 ORF TF | GTGATGGGTCAACCAGAGAG | ΔFvpex13 mutant screen |
| Pex13 ORF TR | CAGGACAAAGAAGAGCAAAG | |
| Pex13 LINK TF | GTCGTCGTTTCCGATTTGTTTT | ΔFvpex13 mutant screen |
| HY/R | GATGTAGGAGGGCGTGGATATGTCCT | |
| Pex13 locF | GAACAAAAGCTGGGTCTCCGTGGTTGGGTTTCT | ΔFvpex13 complementation and GFP fusion |
| Pex13 locR | CTGCAGGCATGCAAG CGAGTAGAAGTGGCTCCTCTGCATG | |
| Pex13 locSF | AACCCAATCTTCAAACTCATGGCCTCTCCACCCAAGCC | |
| Pex13 locMF | TCTTCAAAATGGCTCAGTGGCTGATCCCGATTCTA | FvPex13 mCherry fusion |
| Pex13 locSMF | TCTTCAAAATGGCTCATGGCCTCTCCACCCAAGCC | |
| Pex13 locMR | CTTGCTCACCATACTCGAGTAGAAGTGGCTCCTCTG | |
| Pex14 AFF | AGGAGCGGTAGATTGACGAG | ΔFvpex33 deletion and probe, ΔFvpex14/33 probe |
| Pex14 AFR | TTGACCTCCACTAGCTCCAGCCAAGCCTGAGCGATGATGATGGGAAC | |
| Pex14 BFF | GAATAGAGTAGATGCCGACCGCGGGTTCTGTTGAATGGAGGTGTTTA | |
| Pex14 BFR | GTTGCTTTGCCACTTGTAAT | |
| Pex14ORF TF | CTATGAGTTAGGCAAGGTTTGTG | ΔFvpex33 mutant screen |
| Pex14 ORF TR | TTGAGTTGAGGAGCAGAGGG | |
| Pex14 LINK TF | TGATGCGGTCACCACTAACG | ΔFvpex33 mutant screen |
| Pex14 locF | GAACAAAAGCTGGGTCGGCGATTTGAGCACATTTT | ΔFvpex33complementation and GFP fusion |
| Pex14 locR | CTGCAGGCATGCAAG AGAGCTGGAGCTAGCCCCAGCCTTT | |
| Pex14 locMF | TCTTCAAAATGGCTCCGGCGATTTGAGCACATTTT | FvPex14 mCherry fusion |
| Pex14 locMR | CTTGCTCACCATACTAGAGCTGGAGCTAGCCCCAGCCTTT | |
| Pex14 AFR-N: | AATGGAAATTGTAAGCGTTAATCTAGATGAGCGATGATGATGGGAAC | ΔFvpex14 *and* ΔFvpex33 mutant screen |
| Pex14 BFF-N | TATCGCCTTCTTGACGAGTTCTTCTGACTGTTGAATGGAGGTGTTTA | |
| Pex8 AFF | GTCGCAAGGTATCGGTGGTT | ΔFvpex8 deletion and probe |
| Pex8 AFR | CGCATTCGGATTGTATGGTG | |
| Pex8 BFF | TTGGGTATGTCTCCGTTGAA | |
| Pex8 BFR | CCTGGACGAGCTGACCTGTA | |
| Pex8 ORF TF | TGGCGGTTGAATCGTTTGAG | ΔFvpex8 mutant screen |
| Pex8 ORF TR | TCGGTTGGCAGAGTGAGTGG | |
| Pex8 LINK TF | CTAAAGCGAGAACGTAGCCC | ΔFvpex8 mutant screen |
| Pex8 locF | GAACAAAAGCTGGGTTTGTCTGCGGTGGTGTCA | ΔFvpex8 complementation and GFP fusion |
| Pex8 locR | CTGCAGGCATGCAAGCAGATGGCTAGACCTGTCAG | |
| HYG/F | GGCTTGGCTGGAGCTAGTGGAGGTCAA | Gene replacement by *HPH* |
| HY/R | GTATTGACCGATTCCTTGCGGTCCGAA | |
| YG/F | GATGTAGGAGGGCGTGGATATGTCCT | |
| HYG/R | AACCCGCGGTCGGCATCTACTCTATTC | |
| G418F1 | TCTAGATTAACGCTTACAATTTCCA | Gene replacement by *NEO* |

| | | |
|---|---|---|
| G418R1 | GCCCAATAGCAGCCAGTCC | |
| G418F2 | CAACAACACGCATCATCCCA | |
| G418R2 | TCAGAAGAACTCGTCAAGAA | |
| Pex13 BDF | GGAATTCCATATGATGGCCTCTCCACCCAAGCC | Y2H Pex13 AD vector<br>Y2H Pex13 BD vector |
| Pex13 BDR: | CGGGATCCCTACGAGTAGAAGTGGCTCCTCTG | |
| Pex14 BDF : | GGAATTCCATATGATGGCTATTCGCGAGGATCTCG | Y2H Pex14 AD vector<br>Y2H Pex14 BD vector |
| Pex14 BDR: | ACGGGATCCTCAAGAGCTGGAGCTAGCCCC | |
| BDF Pex8 | GCCATGGAGGCCGAAATGTCCTCCGATAGACTCCT | Y2H Pex8 BD vector |
| BDR Pex8 | CTGCAGGTCGACGGATTACAGATGGCTAGACCTGT | |
| BDF Pex33 | GCCATGGAGGCCGAAATGAGCGACTCAGATTCCAA | Y2H Pex33 BD vector |
| BDR Pex33 | CTGCAGGTCGACGGATCATCTTGCCGAGGCGGGGA | |
| Pex33 ADF: | GGAATTCCATATGATGAGCGACTCAGATTCCAA | Y2H Pex33 AD vector |
| Pex33 ADR: | CGGGATCCTCATCTTGCCGAGGCGGGGA | |
| Pex8 ADF: | GGAATTCCATATGATGTCCTCCGATAGACTCCT | Y2H Pex8 AD vector |
| Pex8 ADR: | TCCCCCCGGGTTACAGATGGCTAGACCTGT | |
| Pex5 ADF: | TCCCCCGGGAATGTCGTTTATGGGTGGCGCT | Y2H Pex5 AD vector |
| Pex5 ADR: | CCATCGATCTAAAACTCAAATTCCGGTCTGAA | |
| Pex7 ADF: | GGAATTCCATATGATGGCTGCAATGCTTGAATT | Y2H Pex7 AD vector |
| Pex7 ADR: | CGGGATCCTCATGGTATTCGTTGGCCCA | |
| Tubulin QF | TCTGACTTCAGGAATGGTCGTTAC | qRT-PCR |
| Tubulin QR | AGCGGTCTGGATGTTGTTGG | |
| FUM1QF | GTTCAGGGCAAGTCCATCATTA | qRT-PCR |
| FUM1QR | CGGTATCGCCATCATCGTC | |
| FUM8QF | TCGGATTGCCACGGTCAC | qRT-PCR |
| FUM8QR | GCACGGTTCCAAACTTCAGG | |
| FUM19QF | GTTCCTCCTGTAGCCATT | qRT-PCR |
| FUM19QR | GCGTCTAATAGCCACCAT | |
| FUM21QF | TCCTGTTCAGAAGAATAC | qRT-PCR |
| FUM21QR | CTCAGTAACCTCATCTAC | |
| PEX1 QF | CGACAAGATTCATTATTCCATAC | qRT-PCR |
| PEX1 QR | ACAAGCGGTTCAAAGTTT | |
| PEX2 QF | AATATACCGACGCCAGAA | qRT-PCR |
| PEX2 QR | TCCAAGCATAGTTTCCAAAT | |
| PEX5 QF | ATAAGGCTAAGGAACAAGAAG | qRT-PCR |
| PEX5 QR | GTTGCTCCATTCGGTATC | |
| PEX7 QF | TCTCATAAGTGCCAGTTA | qRT-PCR |
| PEX7 QR | ACGATTCATAATTCCCATT | |
| PEX8 QF | TAACATCTCTGAACACTTA | qRT-PCR |
| PEX8 QR | GATATGGTTGATAGAATAGC | |
| PEX13 QF | GATGGTAACCCACTTCCT | qRT-PCR |
| PEX13 QR | AGCAAGCGTCCTTATCAT | |
| PEX14 QF | GGAGAAATTGGAACAAGATAAG | qRT-PCR |
| PEX14 QR | TTCGGCATTCTTCAGACT | |

| | | |
|---|---|---|
| 02652qf2 | CTCATCTCACAGTCTCAT | qRT-PCR |
| 02652qR2 | ATCATCAGTAGCAGTATCT | |
| PEX33 QF | ACAGACTCCAAGAATAAC | qRT-PCR |
| PEX33QR | GCAGTATATGTCATTGTG | |
| PTS2 locF | GAACAAAAGCTGGGTAAAGAGTGGAATAGGTCGCA | PTS2 GFP fusion |
| PTS2 locR | CTGCAGGCATGCAAGGACCTGCTCGTTGACAAACA | |
| PTS1 locF3 | GGGTACCGGGCCCCCCCTCGAGGATGGGAAAATGCTTCGA | PTS1 GFP fusion |
| PTS1 PR3 | TCCTCGCCCTTGCTCACCATGGCCAACGCGAAAGACAAGC | |
| GFPF | ATGGTGAGCAAGGGCGAGGA | |
| GFPR | AAGATCTACCATGTACAGCT | |
| PTS1GF3 | CACTCACGGCATGGACGAGCTGTACAAGATGTTGTTGACTTCAGCCCG | |
| PTS1 locR3 | CCCCCGGGCTGCAGGAATTCTTACAGTTTAGCCTTGGGGG | |
| PEX7 locF | GAACAAAAGCTGGGTCTATGCCAGCCGACATTC | GFP fusion |
| PEX7 locR | CTGCAGGCATGCAAGTGGTATTCGTTGGCCCATCA | |
| PEX5 locF | GAACAAAAGCTGGGTCATCTTCGGCATCCTTTC | GFP fusion |
| PEX5 locR | CTGCAGGCATGCAAGAAACTCAAATTCCGGTCTGAACACA | GFP fusion |
| PEX5 locMF | TCTTCAAAATGGCTCCATCTTCGGCATCCTTTC | mCherry fusion |
| PEX5 locMR | CTTGCTCACCATACTAAACTCAAATTCCGGTCTGAACACA | |
| PEX19 LocF | GAACAAAAGCTGGGTGTTTCCGTAATGCGTGGAGT | GFP fusion |
| PEX19 LocR | CTGCAGGCATGCAAGCTGAGGGTTACAGGCTTCATCTCCT | |
| PEX33 locF2 | GAACAAAAGCTGGGTTGGTGGGTGTTGTTTTGCTG | GFP fusion |
| PEX33 locR2 | CTGCAGGCATGCAAGTCTTGCCGAGGCGGGGAAGTTTCTC | |
